# The autism-associated gene SYNGAP1 regulates human cortical neurogenesis

**DOI:** 10.1101/2022.05.10.491244

**Authors:** Marcella Birtele, Ashley Del Dosso, Tiantian Xu, Tuan Nguyen, Brent Wilkinson, Negar Hosseini, Sarah Nguyen, Jean-Paul Urenda, Gavin Knight, Camilo Rojas, Ilse Flores, Alexander Atamian, Roger Moore, Ritin Sharma, Patrick Pirrotte, Randolph S. Ashton, Eric J. Huang, Gavin Rumbaugh, Marcelo P. Coba, Giorgia Quadrato

## Abstract

Autism spectrum disorder (ASD) is a genetically heterogeneous disorder linked with rare, inherited and *de novo* mutations occurring in two main functional gene categories: gene expression regulation and synaptic function^1^. Accumulating evidence points to dysregulation in cortical neurogenesis as a convergent mechanism in ASD pathophysiology^2-8^. While asynchronous development has been identified as a shared feature among ASD-risk genes in the category of gene expression regulation, it remains unknown whether this phenotype is also associated with ASD-risk genes in the synaptic function category. Here we show for the first time the expression of the synaptic Ras GTP-ase activating protein 1 (SYNGAP1), one of the top ASD risk genes^9^, in human cortical progenitors (hCPs). Interestingly, we found that multiple components of the postsynaptic density (PSD) of excitatory synapses, of which SYNGAP1 is one of the most abundant components ^10,11^, are enriched in the proteome of hCPs. Specifically, we discover that SYNGAP1 is expressed within the apical domain of human radial glia cells (hRGCs) where it lines the wall of the developing cortical ventricular zone colocalizing with the tight junction-associated protein and MAGUK family member TJP1. In a cortical organoid model of SYNGAP1 haploinsufficiency, we show dysregulated cytoskeletal dynamics that impair the scaffolding and division plane of hRGCs, resulting in disrupted lamination of the cortical plate and accelerated maturation of cortical projection neurons. Overall, the discovery of the expression and function of SYNGAP1 in cortical progenitor cells reframes our understanding of the pathophysiology of SYNGAP1-related disorders and, more broadly, underscores the importance of dissecting the role of synaptic genes associated with neurodevelopmental disorders in distinct cell types across developmental stages.

## Introduction

Exome sequencing analyses have identified two major functional categories of genes associated with Autism Spectrum Disorder (ASD): gene expression regulation and synaptic function^1^. SYNGAP1 is a top ASD genetic risk factor ^9^ and one of the most abundant proteins found at the postsynaptic density (PSD) of excitatory synapses^10,11^. Within the PSD, SYNGAP1 functions as a RAS GTPase-activating (RASGAP) protein that regulates synaptic plasticity ^10-15^. Through its RASGAP domain, SYNGAP1 limits the activity of the mitogen-activated protein kinase 1 (Mapk1/Erk2), whereas through its PDZ-binding domain, SYNGAP1 helps assemble the core scaffold machinery of the PSD^16-19^. Despite its classification as a synaptic protein, several lines of evidence suggest a potential role for SYNGAP1 at early stages of cortical neurogenesis. First, homozygous deletion of Syngap1 in embryonic mice leads to early developmental lethality ^20^. Second, decreased *syngap1* levels have been shown to affect the ratios of neural progenitor cells to mature neurons in Xenopus tropicalis ^21^. Third, disruption of the Syngap1 signaling complex in embryonic mice results in deficits in the tangential migration of GABAergic interneurons ^22^.Lastly, in addition to being an ASD genetic risk factor, *de novo* mutations in SYNGAP1 have been found in patients with intellectual disability, epilepsy, neurodevelopmental disability, and global developmental delay ^23-25^. This evidence, combined with the high frequency and penetrance of pathogenic SYNGAP1 variants, indicates a major and unique role for SYNGAP1 in human brain development. However, as with other components of the scaffold machinery of the PSD, it remains unclear if SYNGAP1 is expressed in early cortical progenitors and how it affects cortical neurogenesis.

To address these questions, we used 3D cultures of human brain organoids. Derived from human embryonic or induced pluripotent stem cells (hPSCs), organoids have emerged as an effective way to model genetic architecture and cellular features of human brain development and disease ^4,5,26-33^. These reproducible models of the human forebrain are capable of generating cellular diversity and epigenetic states that follow the developmental trajectory of the corresponding endogenous cell types ^2,34-38^, allowing for the functional characterization of ASD-risk genes in a longitudinal modeling and human cellular context.

Here, we show for the first time the expression of SYNGAP1 protein in human radial glia cells (hRGCs). Mechanistically, we find that SYNGAP1 regulates cytoskeletal remodeling of subcellular and intercellular components of hRGCs, with haploinsufficiency leading to disrupted organization of the developing cortical plate. By performing single-cell transcriptomics coupled with structural and functional analysis of mutant organoids, we discovered that SYNGAP1 regulates the timing of hRGCs differentiation with haploinsufficient organoids exhibiting accelerated maturation of cortical projection neurons. Additionally, we show that reduction in SynGAP1 levels affects cortical neurogenesis in a murine model suggesting that SynGAP1 function in cortical progenitors is conserved across species. Altogether, these findings reveal a novel function for the classically defined synaptic protein SYNGAP1 at early stages of human cortical neurogenesis, providing a new framework for understanding ASD pathophysiology.

## Results

### SYNGAP1 is expressed in human ventricular radial glia and colocalizes with the tight junction protein TJP1

Syngap1 expression has been reported to be largely restricted to the PSD of mature excitatory synapses^10,11^ of mouse cortical projection neurons, where it has a cell type-specific function^39,40^. To investigate the expression of SYNGAP1 in distinct human cortical cell types, we took advantage of a recently published single cell RNA-seq data set of early human fetal telencephalic/cortical development from post-conception days (PCD) 26 to 54 ^41^. We compared SYNGAP1 expression and distribution across all developmental stages with the expression of well-known gene markers for neuroepithelial/radial glial cells (PAX6), radial glial cells (HES5), and intermediate progenitor cells (IPC) (EOMES/TBR2). Within the data set, SYNGAP1 expression was detected throughout the age range. Interestingly, SYNGAP1’s expression levels were enriched in hRGCs (Suppl. Fig 1a-c), pointing to a novel function of SYNGAP1 in human cortical progenitors.

To further validate SYNGAP1 protein expression at this stage of development, we performed a proteome profiling of human cortical organoids (D.I.V. 7), which are entirely composed of SOX2+ and PAX6+ progenitors at this stage of development (Suppl. Fig. 1d). Using this pipeline, we were able to identify a total of 8690 proteins (Suppl. Table 1), including 24 unique peptides for SYNGAP1. Interestingly, 793 of these proteins belonged to the SynGO ontology term ‘Synapse’(Suppl. Table 2), with enrichment in components of the postsynaptic density, pointing to an unappreciated role of classically defined postsynaptic proteins in progenitor biology (Fig. 1a). Within 2-month-old organoids, the Ventricular Zone (VZ) is delineated by radially aligned SOX2+ neural stem cells surrounding the apical junctional belt, which is composed of adherens and tight junction proteins such as TJP1. Immunofluorescence using a SYNGAP1 antibody (Suppl. Fig 1f-g) revealed that SYNGAP1 expression was enriched at the apical end feet of the VZ cells (Fig. 1b) as well as in MAP2+ neurons (Fig.1d). Notably, SYNGAP1 and TJP1 co-localize (Fig. 1b). Through a series of immunohistochemical analyses of micro-dissected VZ and SVZ specimens of human cortex at gestational week 17 (GW17) (Fig. 1f) and mouse cortex at E13.5 (Suppl. Fig. 1e), we confirmed the same pattern of SYNGAP1 expression in the apical end feet of ventricular radial glia cells. Moreover, SYNGAP1 immunoprecipitation and mass spectrometry analysis shows that TJP1 is part of the SYNGAP1 interactome (Fig. 1c, Suppl. Table 3). The PDZ ligand domain of SYNGAP1 is known to mediate localization of the protein to the PSD via binding to MAGUK proteins such as Dlg4 in mature rodent neurons^42^.TJP1 is a MAGUK family member and shares the modular-domain composition of other MAGUK proteins, and therefore can associate to SYNGAP1 through a PDZ domain. Therefore, we hypothesize that SYNGAP1 interacts with TJP1, a PDZ domain containing protein, in a similar manner (Fig. 1e). The capacity of *SYNGAP1* to associate to proteins containing PDZ domains is isoform-dependent^16,43^. To address whether hRGCs express the SYNGAP1 isoform alpha 1 (id: A0A2R8Y6T2), the isoform with the capacity to associate to PDZ domains, we designed a parallel reaction monitoring experiment based on high resolution and high precision mass spectrometry. This assay allowed us to show that similarly to what occurs in mature rodent synapses^16,43^, SYNGAP1 has the capacity to associate to PDZ containing proteins in hRGCs cells (Fig. 1e, Suppl. Fig. 2a-b).

**Figure 1.**
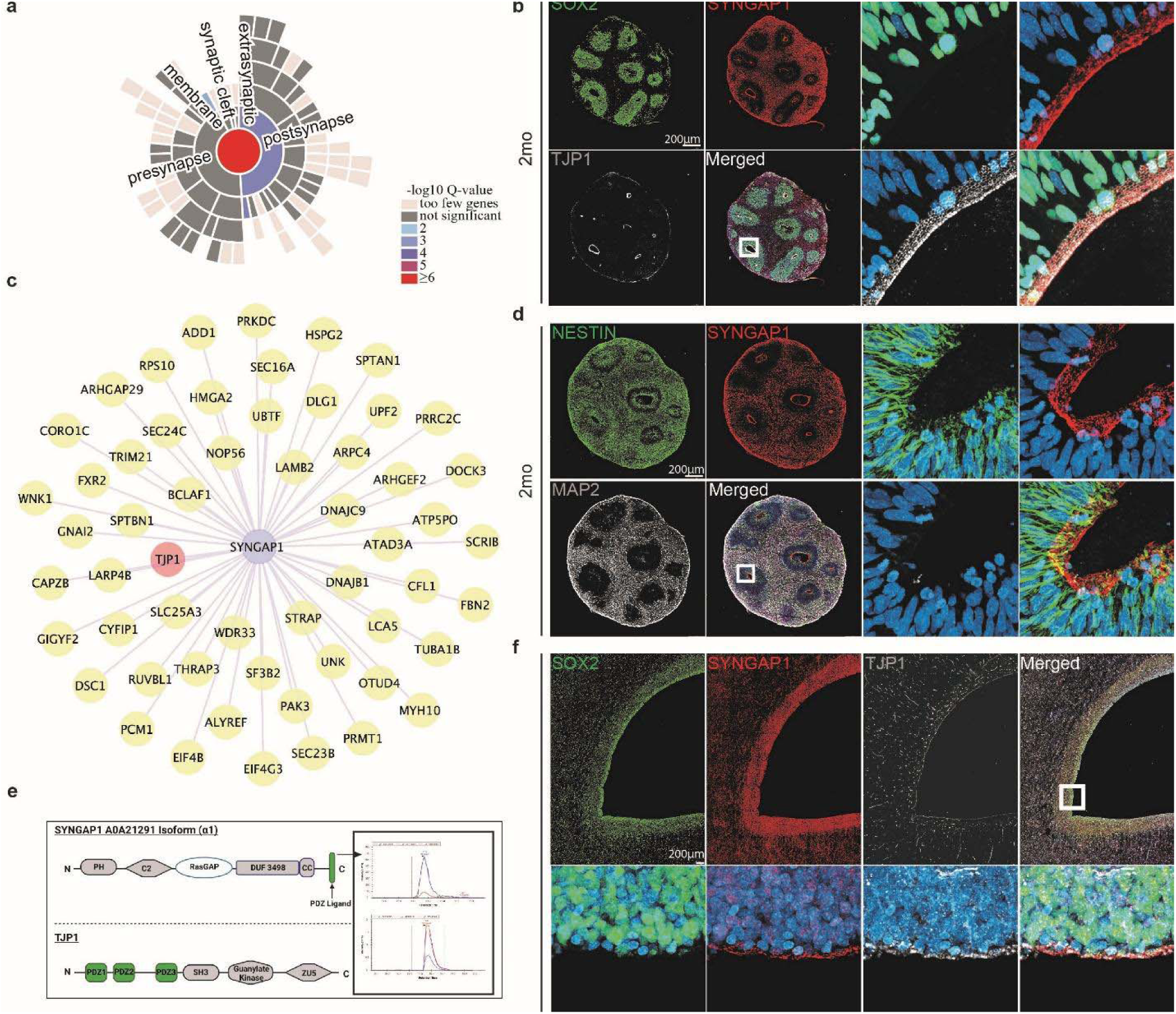
SYNGAP1 is expressed in human radial glia progenitors and colocalizes with the tight junction protein TJP1. A. SynGO analysis results from D.I.V. 7 corrected organoids proteomic data set. 21 organoids from 3 independent experiments were analyzed. B. Two-month-old control organoid stained for the neural progenitor marker SOX2, the tight junction protein TJP1 and SYNGAP1. SYNGAP1 is highly expressed at the ventricular wall. White box indicates the region of interest selected for the merged images showing colocalization of the nuclear marker DAPI, TJP1, and SYNGAP1. C. Schematic for the protein interaction network of *SYNGAP1*. Analysis performed on 21 organoids from 3 independent experiments at D.I.V. 7. The tight junction protein TJP1 is highlighted in pink. D. Two-month-old Patient^Corrected^ organoid stained for the radial glial marker NESTIN, the neuronal marker MAP2, and SYNGAP1. SYNGAP1 is highly expressed within mature MAP2 positive neuronal populations outside of the VZ, as well as in NESTIN positive cells at the ventricle wall. White box indicates the region of interest selected for the merged images showing colocalization of DAPI, SYNGAP1, and NESTIN positive cells. E. Schematic of key functional domains within the SYNGAP1 alpha 1 isoform and TJP1 proteins including representative spectra of the two identified peptides for the alpha1 isoform. F. Immunohistochemical staining of the human brain at gestational week 17. Tissue section is from the prefrontal cortex at the level of the lateral ventricle and medial ganglionic eminence. White box indicates the region of interest selected for the merged images showing colocalization of DAPI, TJP1, and SYNGAP1.

### SYNGAP1 plays a role in the cytoskeletal organization of hRGCs

To determine the function of SYNGAP1 in hRGCs, we developed an early cortical organoid model (D.I.V. 7) from a patient carrying a SYNGAP1 truncating (p.Q503X) mutation. This patient presented with intellectual disability, developmental delay, autistic features and epilepsy. For this, we generated iPSC lines from the patient and the correspondent corrected isogenic control (Suppl. Fig. 3b,c,g,h Suppl. Table 4-5). The characterization of the patient-derived iPSC cell line shows that the p.Q503X mutation produces a haploinsufficent model of SYNGAP1 disfunction with a 51.6% decrease in SYNGAP1 total protein levels and not detectable protein fragments as evidenced by WB and quantitative Mass Spectrometry assays (Suppl. Fig. 3d-f) To start to address the role of SYNGAP1 in hRGCs we performed bulk RNA sequencing of 7 D.I.V. organoids from the Patient ^p.Q503X^ and Patient^Corrected^ lines, and our Gene Ontology (GO) analyses found terms for biological processes related to cytoskeletal remodeling and migration to be significantly enriched within our gene list (P value = 1.02e-03) (Fig. 2a; Suppl. Table 6). In addition, GO analysis of the SYNGAP1 interactome revealed that the predicted set of molecular functions regulated by SYNGAP1 clustered within the functional categories of cytoskeletal organization and regulation (Suppl. Fig. 3a).

**Figure 2.**
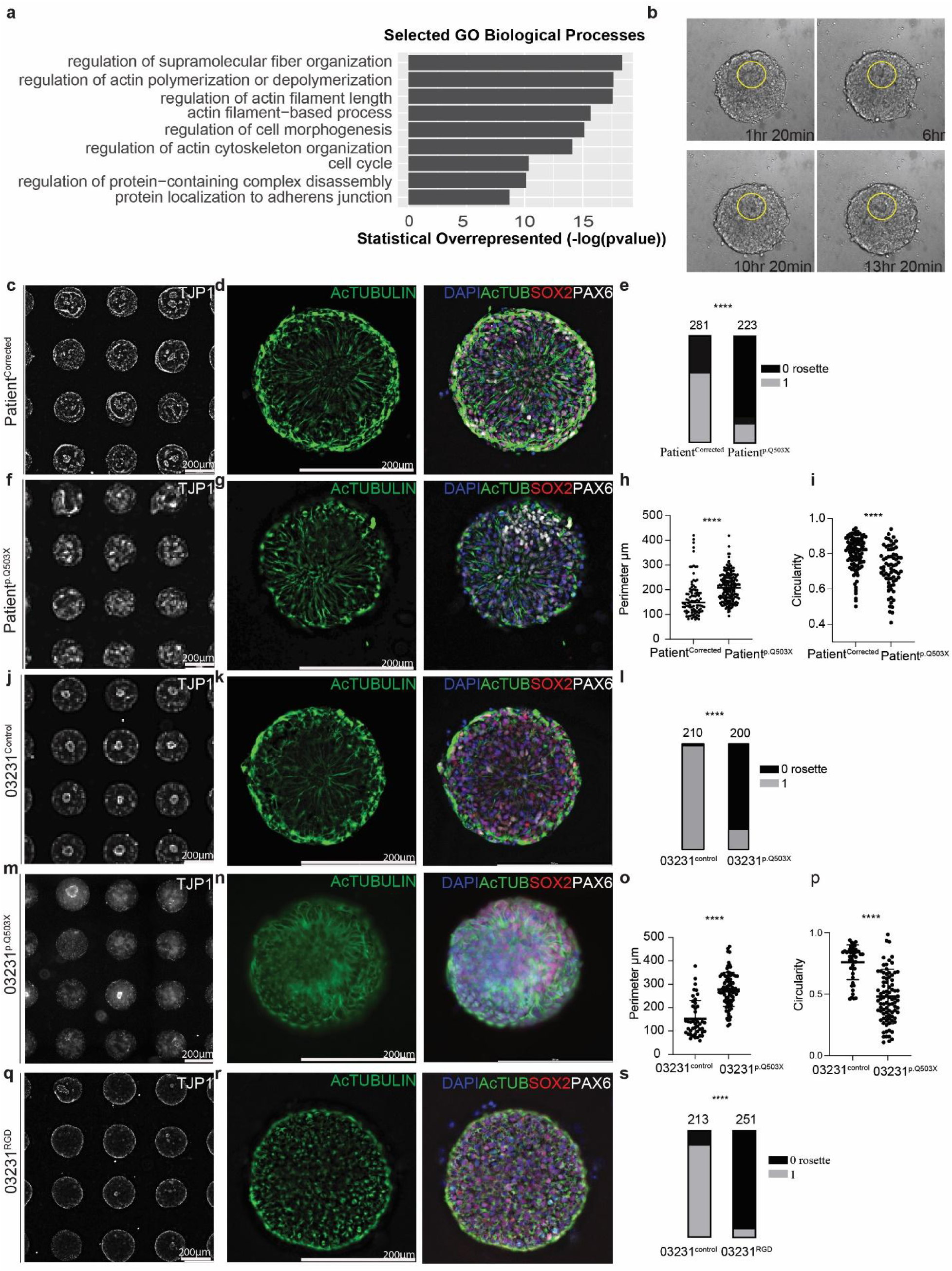
SYNGAP1 plays a role in the cytoskeletal organization of human radial glia. A. Selected GO terms for biological processes for bulk RNA sequencing data collected from D.I.V. 7 control organoids. A total of 100 organoids from 2 independent experiments were analyzed. B. Still frames taken from live imaging of Patient^Corrected^ rosette formation from day 5 to day 7 after initial seeding. C. Array of single rosettes generated from the Patient^Corrected^ line control labeled with TJP1 staining. D. A single rosette generated from Patient^Corrected^ iPSCs. The rosette is composed of cells positive for the neural stem cell marker SOX2 and radial glial progenitors positive for PAX6. Acetylated tubulin labels the microtubules of radial glial progenitors and highlights their radial organization around the luminal space. E. Quantification of the number of rosettes successfully formed in the Patient^Corrected^ and Patient^p.Q503X^ lines. Successful rosette formation was defined as containing a densely packed central area positive for TJP1 and radial organization of surrounding progenitor cells. Chi-square test was performed on n=281 control single rosettes and n= 223 Patient^p.Q503X^ single rosettes from 3 independent experiments. P value <0.0001. F. Array of single rosettes generated from the Patient^p.Q503X^ line labeled with TJP1 staining. G. A single rosette generated from Patient^p.Q503X^ iPSCs. Similar to the Patient^Corrected^ rosette, it is composed of cells positive for SOX2 and PAX6. Acetylated tubulin labels the microtubules of radial glial progenitors and shows diminished radial organization around the luminal space as compared to the Patient^Corrected^ rosette. H. Quantification of the rosette lumen perimeter, as marked by TJP1 labeling in the Patient^Corrected^ and Patient^p.Q503X^ lines. Unpaired t-test was performed on n=94 corrected single rosettes and n= 198 Patient^p.Q503X^ single rosettes from 3 independent experiments. P value <0.0001. I. Quantification of the circularity of the rosette lumen as marked by TJP1 labeling in the Patient^Corrected^ and Patient^p.Q503X^ lines. Unpaired t-test was performed on n=99 corrected single rosettes and n= 69 Patient^p.Q503X^ single rosettes from 3 independent experiments. P value <0.0001. J. Array of single rosettes generated from the 03231^Control^ line labeled with TJP1 staining. K. A single rosette generated from 03231^Control^ iPSCs. It is composed of cells positive for SOX2 and PAX6. Acetylated tubulin labels the microtubules of radial glial progenitors and shows radial organization and central luminal space. L. Quantification of the number of rosettes successfully formed in the 03231^Control^ and 03231^p.Q503X^ lines. Successful rosette formation was defined as containing a densely packed central area positive for TJP1 and radial organization of surrounding progenitor cells. Chi-square test was performed on n=210 03231^Control^ single rosettes and n= 200 03231^p.Q503X^ single rosettes. P value <0.0001. Analysis was performed from 3 independent experiments. M. Array of single rosettes generated from the 03231^p.Q503X^ line labeled with TJP1 staining. N. A single rosette generated from 03231^p.Q503X^ iPSCs. Similar to the 03231^Control^ rosette, it is composed of cells positive for SOX2 and PAX6. Acetylated tubulin labels the microtubules of radial glial progenitors and shows little to no radial organization around the luminal space as compared to the 03231^Control^ rosette. O. Quantification of the perimeter of the rosette lumen as marked by TJP1 labeling in the 03231^Control^ and 03231^p.Q503X^ lines. Unpaired t-test was performed on n=45 03231^Control^ single rosettes and n= 88 03231^p.Q503X^ single rosettes from 3 independent experiments. P value <0.0001. P. Quantification of the circularity of the rosette lumen as marked by TJP1 labeling in the 03231^control^ and 03231^p.Q503X^ lines. Unpaired t-test was performed on n=45 03231^Control^ single rosettes and n= 88 03231^p.Q503X^ single rosettes from 3 independent experiments. P value <0.0001 Q. Array of single rosettes generated from the 03231^RGD^ line labeled with TJP1 staining. R. A single rosette generated from 03231^RGD^ iPSCs. Similar to the 03231^Control^ rosette, it is composed of cells positive for SOX2 and PAX6. Acetylated tubulin labels the microtubules of radial glial progenitors and shows little to no radial organization or central luminal space. S. Quantification of the number of rosettes successfully formed in the 03231^Control^ and 03231^RGD^ lines. Successful rosette formation was defined as containing a densely packed central area positive for TJP1 and radial organization of surrounding progenitor cells. Chi-square test was performed on n=213 03231^Control^ single rosettes and n= 251 03231^RGD^ single rosettes. P value <0.0001. Analysis was performed from 3 independent experiments.

hRGCs have a bipolar shape with distinct apical and basolateral domains and include a process terminating at the ventricular surface and another process reaching the pial surface. To monitor the effect of SYNGAP1 haploinsufficiency on the shape and polarity of hRGCs processes, we employed a standardized single neural rosette protocol^44^ that allows high-throughput generation and analysis of rosette formation (Fig. 2b, Suppl. Video 1). We found homogeneous expression of the cortical progenitor markers PAX6 and SOX2 with radially organized acetylated tubulin networks in rosettes derived from the Patient^Corrected^ line (Fig. 2c,d, Suppl. Fig. 4a). However, these structures were less organized in SYNGAP1 haploinsufficient tissues, and their formation was less frequent than in corrected rosettes (Fig. 2e-h, Suppl. Fig. 4b). Through a high throughput analysis of the localization tight-junction marker TJP1 in single rosettes tissues, we found that SYNGAP1 haploinsufficient rosettes exhibited apico-basal polarity from a wider, more irregular TJP1 positive region, while corrected individual rosettes had a tighter and more circular TJP1 ring (Fig. 2i, Suppl. Fig. 4d-e).

To understand the contribution of the patient’s genetic background to the observed phenotypes, we repeated this assay in rosettes derived from a new line that we generated to carry the truncating p.Q503X mutation in the control 03231 background ^45^ (Suppl. Fig. 3 j, m-o). Importantly, single rosettes generated from this line displayed the same phenotypes as Patient^p.Q503X^ (Fig. 2j-p; Suppl. Fig. 4c,f,g,i). This indicates that the SYNGAP1 haploinsufficiency phenotype is not influenced by the individual genomic context. This finding is consistent with the high penetrance of pathogenic SYNGAP1 variants in patients. The ability of SYNGAP1 to partake in cytoskeletal remodeling in mature neurons has been attributed to enzymatic activity from its RAS GTPase-activating domain^10,11,40,46,47^. To assess if SYNGAP1’s enzymatic function is conserved in hRGCs, we edited the 03231 iPSC line to carry a homozygous non-functional RASGAP domain. We named this RASGAP-Dead line 03231^RGD^ (Suppl. Fig. 3 i-l; Suppl. Table 4) and compared it to its isogenic control iPSC line. We observed a more marked effect compared to the SYNGAP1 haploinsufficient phenotype, with the tissue generated from the RASGAP-Dead (RGD) line displaying fully disrupted apico-basal polarity and a reduced central TJP1 positive luminal space (Fig. 2q-s; Suppl. Fig.4h,j). Importantly, these profound defects lead to the loss of about 90% of the 03231^RGD^ organoids over 2 months preventing a thorough characterization of organoids at later stages of development (Suppl. Fig. 4k). These data indicate that impairment in the cytoskeletal architecture of apical end feet, as well as disruption of radial elongation of the basal process of hRGCs are present in both SYNGAP1 haploinsufficient and RGD-derived tissues, indicating that this function is dependent on the RASGAP domain of SYNGAP1. However, we cannot exclude that misfolding effects or changes in SYNGAP1 protein localization and interaction occur following mutation of the RASGAP domain in the RGD line, affecting the function of other functional sites in the SYNGAP1 protein. Further experiments will need to be performed to prove the necessity of the enzymatic function of the RASGAP domain in regulating cytoskeleton dynamics of hRGCs.

Altogether, these data show that SYNGAP1 regulates a dynamic cytoskeletal network that influences the subcellular and intercellular organization of ventricular radial glia.

### SYNGAP1 haploinsufficiency disrupts the organization of the developing cortical plate

hRGCs control the generation and organization of a proper VZ by tightly connecting to each other via an AJ belt at their apical endfeet^48^; through their basal process, they guide newly born neurons across the entire thickness of the developing cortex, serving as a central organizer for the assembly of cortical neuronal columns, layers, and circuitry^49-51^. To assess SYNGAP1’s function during cortical plate formation, we analyzed 2-month-old haploinsufficient and corrected cortical organoids. At this stage, organoids form robust VZs and the surrounding regions are populated with both upper and deep layer cortical projection neurons^35^. The VZs, which are defined by highly dense SOX2 positive areas, serve as the germinal niche for the organoids; these regions were significantly reduced in size, number, and organization in SYNGAP1 haploinsufficient organoids derived from two independent genetic backgrounds (Patient^p.Q503X^ and 03231^p.Q503X^) (Fig. 3a-h). The sizable presence of MAP2+ cells within the VZ of haploinsufficient organoids indicates a potentially impaired ability of neuroblasts to migrate away from the ventricular zone towards the cortical plate or an imbalance in direct neurogenesis (Fig. 3i-n). This is also consistent with the disorganization of radial glial progenitors seen in SYNGAP1 haploinsufficient rosettes, which are known to serve as migratory scaffolding for neuroblasts *in vivo*. Consistent with these results, binning analysis of BrdU labeling experiments in 2-month-old organoids treated for 2 hours with BrdU followed by 24 hours chase revealed the presence of a higher number of BrdU+/ NeuN+ neurons in bins closer to the ventricular wall in SYNGAP1 haploinsufficient organoids (Fig. 3o-t, Suppl. Fig. 5a-d). This further highlights the role of SYNGAP1 in controlling neuronal positioning and the overall organization of the cortical plate during human neurogenesis. In addition, we observed that dividing cells, in haploinsufficient organoids, have a higher propensity to differentiate into NeuN+ neurons while control organoids maintained a larger BrdU+/SOX2+ progenitor pool (Fig. o, r-t, Suppl. Fig. a-d), suggesting a potential impairment in the division mode of hRGCs.

**Figure 3.**
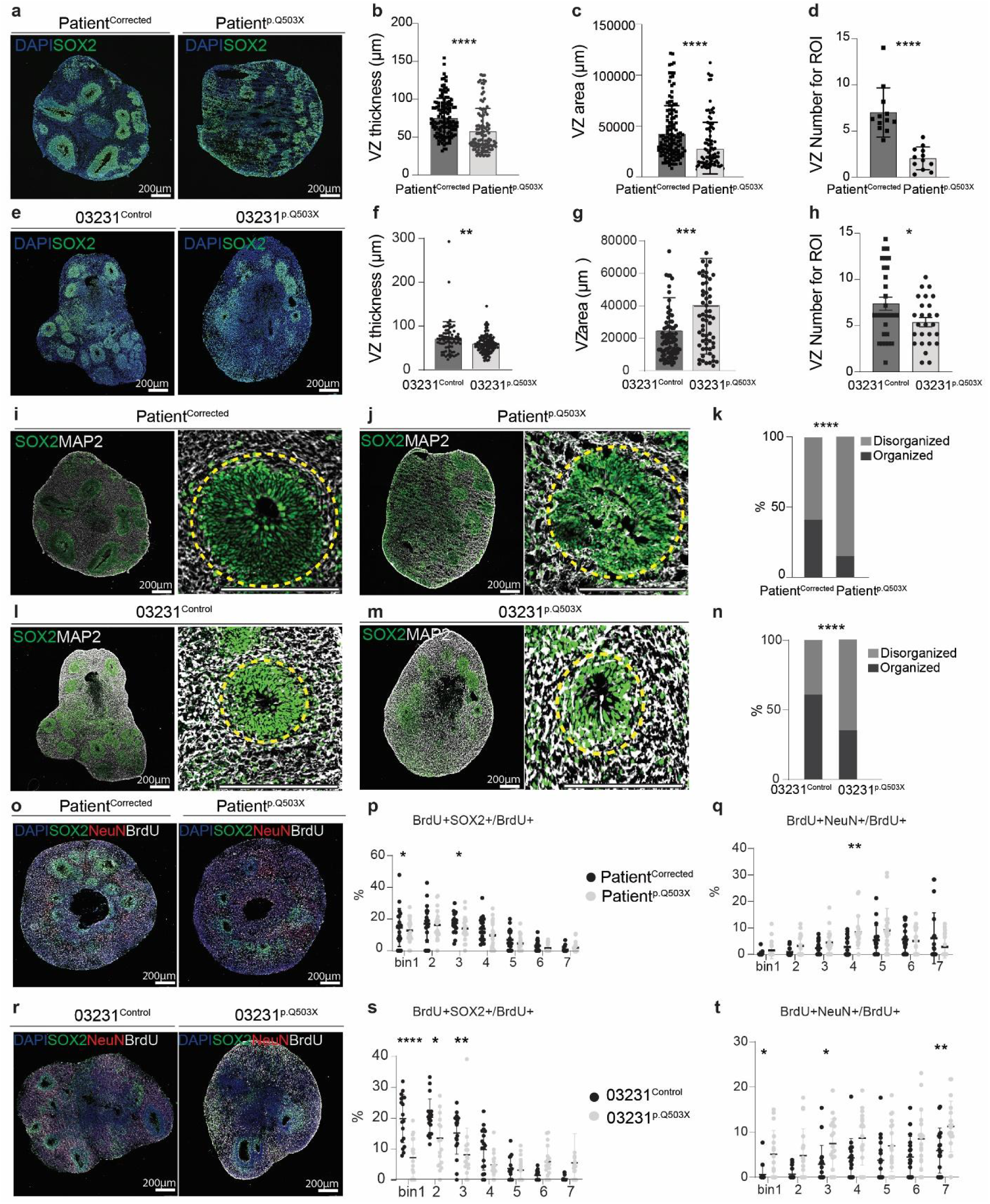
SYNGAP1 haploinsufficiency disrupts the organization of the developing cortical plate. A. Two-month-old organoids show expression of SOX2, a neural stem cell marker, and nuclei marker DAPI. Patient^Corrected^ organoids display SOX2 expression exclusively within the ventricular zone (VZ), while Patient^p.Q503X^ organoids lack a defined VZ. B. VZ analysis shows VZ thickness (µm) in Patient^p.Q503X^ and Patient^corrected^ organoids. Patient^p.Q503X^ organoids present less proliferative VZ compared to Patient^corrected^. Student’s t-test was performed on 4 independent experiments where 20-30 VZs from 3 organoids were analyzed in each experiment. Single dots represent individual VZs. P value (<0.0001). Data is shown as mean ± SD. C. VZ analysis shows VZ area (µm^2^) in Patient^p.Q503X^ and Patient^Corrected^ organoids. SYNGAP1^p.Q503X^ organoids present less extended VZ compared to Patient^Corrected^. Student’s t-test was performed on 4 independent experiments where 20-30 VZs from 3 organoids were analyzed in each experiment. Single dots represent individual VZs. P value (<0.0001). Data is shown as mean ± SD. D. VZ analysis shows VZ number per Region of Interest (ROI) in Patient^p.Q503X^ and Patient^Corrected^ organoids. Patient^p.Q503X^ organoids present fewer VZs compared to corrected. Student’s t-test was performed on 4 independent experiments where 3-6 ROI from 3 organoids were analyzed in each experiment. Single dots represent the mean number of VZs per ROI in individual organoids. P value (<0.0001). Data is shown as mean ± SD. E. Two-month-old organoids show expression of SOX2, a neural stem cell marker, and nuclei marker DAPI. 03231^Control^ organoids display SOX2 expression exclusively within the ventricular zone (VZ), while 03231^p.Q503X^ organoids lack a defined VZ. F. VZ analysis shows VZ thickness (µm) in 03231^p.Q503X^ and 03231^Control^ organoids. 03231^p.Q503X^ organoids present less proliferative VZ compared to 03231^Control^ Student’s t-test was performed on 4 independent experiments where 20-30 VZs from 3 organoids were analyzed in each experiment. Single dots represent individual VZs. P value <0.0099. Data is shown as mean ± SD. G. VZ analysis shows VZ area (µm^2^) in 03231^p.Q503X^ and 03231^Control^ organoids. 03231^p.Q503X^ organoids present less extended VZ compared to 03231^Control^ Student’s t-test was performed on 4 independent experiments where 20-30 VZs from 3 organoids were analyzed in each experiment. Single dots represent individual VZs. P value (<0.0046). Data is shown as mean ± SD. H. VZ analysis shows VZ number per Region of Interest (ROI) in 03231^p.Q503X^ and Patient^Corrected^ organoids. 03231^p.Q503X^ organoids present fewer VZs compared to corrected. Student’s t-test was performed on 4 independent experiments where 3-6 ROI from 3 organoids were analyzed in each experiment. Single dots represent the mean number of VZs per ROI in individual organoids. P value (<0.0134). Data is shown as mean ± SD. I. Two-month-old organoids show expression of SOX2 and microtubule associated protein-2 (MAP2), a pan-neuronal marker. Patient^Corrected^ organoids display SOX2 expression in the VZ, while MAP2 labels neurons that have migrated away from the VZ and now populate the cortical plate. Dashed yellow circles identify VZ regions. J. Patient^p.Q503X^ organoids display SOX2 expression in the VZ and MAP positive neurons inside and outside of the VZ, indicating impaired migration of neurons away from the VZ. Dashed yellow circles identify VZ regions. K. Quantification of the occurrence of organized versus disorganized VZ structures in Patient^Corrected^ and Patient^p.Q503X^ organoids based on SOX2 and MAP2 staining patterns. Chi-square test was performed on mean values of VZs from 3-9 ROI of 3 organoids, analyzed in 4 independent experiments. P value <0.0001. Data is shown as mean ± SD. L. Two-month-old organoids show expression of SOX2 and microtubule associated protein-2 (MAP2), a pan-neuronal marker. 03231^Control^ organoids display SOX2 expression in the VZ, while MAP2 labels neurons that have migrated away from the VZ and now populate the cortical plate. Dashed yellow circles identify VZ regions. M. 03231^p.Q503X^ organoids display SOX2 expression in the VZ and MAP positive neurons inside and outside of the VZ, indicating impaired migration of neurons away from the VZ. Dashed yellow circles identify VZ regions. N. Quantification of the occurrence of organized versus disorganized VZ structures in 03231^Control^ and 03231^p.Q503X^ organoids based on SOX2 and MAP2 staining patterns. Chi-square test was performed on mean values of VZs from 3-9 ROI of 3 organoids, analyzed in 4 independent experiments. P value <0.0001. Data is shown as mean ± SD. O. Patient^Corrected^ and Patient^p.Q503X^ organoids stained for the progenitor marker SOX2, the neuronal marker NeuN and the proliferative marker BrdU. P. Binning analysis for the number of double positive BrdU and SOX2 cells normalized to the total number of BrdU positive cells. The higher number of double positive cells in earlier bins in the Patient^Corrected^ organoids indicates a propensity for progenitors to remain in a proliferative state when compared to Patient^p.Q503X^. A two-way ANOVA was performed on 15 Patient ^p.Q503X^ ventricles and 12 Patient^Corrected^ from 3 independent experiments. P Value Bin 1 = 0.4, P Value Bin 3 = 0.2. Data is shown as mean ± SD. Q. Binning analysis for the number of double positive BrdU and NeuN cells normalized to the total number of BrdU positive cells. The higher number of double positive cells in the Patient^p.Q503X^ organoids indicates a propensity for progenitors to transition to a more differentiated state when compared to Patient^Corrected^. A two-way ANOVA was performed on 15 Patient ^p.Q503X^ ventricles and 12 Patient^Corrected^ from 3 independent experiments. P Value Bin 4 = 0.0094. Data is shown as mean ± SD. R. 03231^Control^ and 03231t^p.Q503X^ organoids stained for the progenitor marker SOX2, the neuronal marker NeuN and the proliferative marker BrdU. S. Binning analysis for the number of double positive BrdU and SOX2 cells normalized to the total number of BrdU positive cells. The higher number of double positive cells in earlier bins in the 03231^Control^ organoids indicates a propensity for progenitors to remain in a proliferative state when compared to 03231^p.Q503X^. A two-way ANOVA was performed on 18 03231^p.Q503X^ ventricles and 18 03231^Control^ ventricles from 3 independent experiments. P Value Bin 1 = 0.0001>, P Value Bin 2 = 0.0110, P Value Bin 3 = 0.0076. Data is shown as mean ± SD. T. Binning analysis for the number of double positive BrdU and NeuN cells normalized to the total number of BrdU positive cells. The higher number of double positive cells in the 03231^p.Q503X^ organoids indicates a propensity for progenitors to transition to a more differentiated state when compared to 03231^Control^. A two-way ANOVA was performed on 18 03231^p.Q503X^ ventricles and 18 03231^Control^ ventricles from 3 independent experiments. P Value Bin 1 = 0.0448, P Value Bin 3 = 0.0400, P Value Bin 7 = 0.0094. Data is shown as mean ± SD.

### SYNGAP1 haploinsufficiency affects the division mode of human radial glial progenitors

The precise spatiotemporal regulation of RGCs division and differentiation^52,53^ is critical for proper lamination and microcircuit assembly in the developing human cortex. During ventricular RGC division, mitotic spindle orientation determines the cleavage plane and predicts cell fate decisions that result in either symmetric proliferative or asymmetric differentiative divisions^54,55^. To better characterize the observed imbalance in the ratio of progenitors to neurons, we analyzed the division planes of ventricular radial glia and found an increased proportion of cells undergoing differentiative divisions in haploinsufficient organoids (Fig. 4a-e). This data suggests an earlier depletion of the progenitor pool due to premature loss of the proliferative potential of the hRGCs in haploinsufficient organoids derived from two independent genetic backgrounds (Patient^p.Q503X^ and 03231^p.Q503X^).

**Figure 4.**
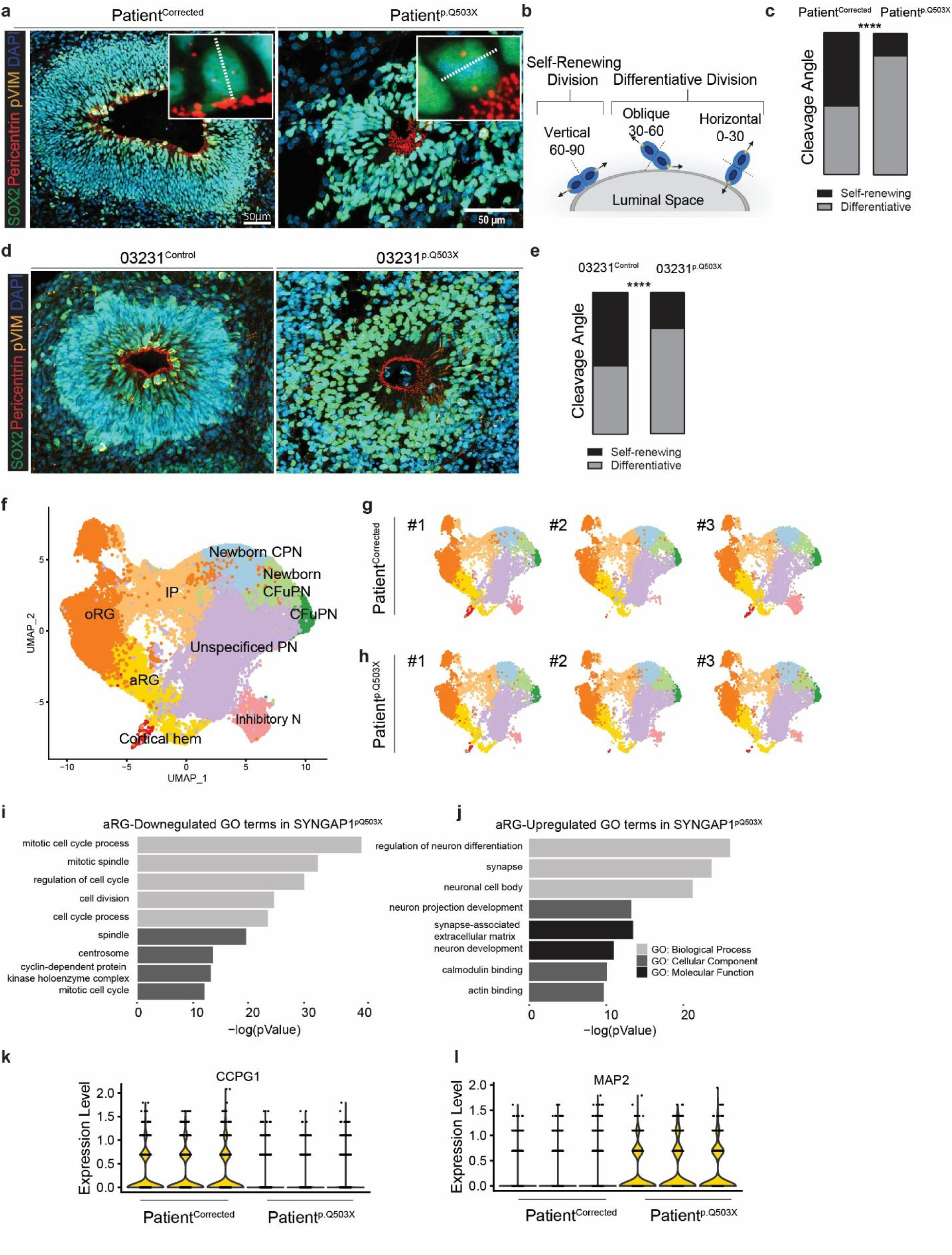
SYNGAP1 haploinsufficiency affects the division mode of human radial Glial progenitors. A. Patient^Corrected^ and Patient^p.Q503X^ organoids display SOX2 expression in the VZ, dividing neural progenitors marked by phospho-vimentin (pVIM), and the centrosome labeling marker Pericentrin. B. Schematic representation of cell divisions at the VZ wall. Schematic illustrates self-renewing divisions (vertical, with angle between 60 to 90 degrees from the apical wall to the mitotic spindle) and differentiative divisions (oblique, 30 to 60 degrees; horizontal, 0 to 30 degrees). C. Cleavage angle analysis showing increased proliferative divisions in cells from Patient^Corrected^ organoids and increased differentiative divisions from Paitent^p.Q503X^ organoids. Mann-Whitney test was performed on a total of 195 cells for each line, from 3 independent experiments. P value < 0.0001. D. 03231^Control^ and 03231^p.Q503X^ organoids display SOX2 expression in the VZ, dividing neural progenitors marked by phospho-vimentin (pVIM), and the centrosome labeling marker Pericentrin. E. Cleavage angle analysis showing increased proliferative divisions in cells from 03231^Control^ organoids and increased differentiative divisions from 03231^p.Q503X^ organoids. Mann-Whitney test was performed on a total of 78 cells for 03231^Control^ and 188 cells 03231^p.Q503X^ cells for condition, from 3 independent experiments. P value < 0.0001. F. Combined t-distributed stochastic neighbor embedding (t-SNE) from single cell RNA sequencing analysis of all organoids at 2 months. G. Individual t-SNE plots for three individual Patient^corrected^ organoids at 2 months. Organoid #1 (n= 8676 cells), Organoid #2 (n= 8377), Organoid #3 (n= 7687). H. Individual t-SNE plots for three individual Patient^p.Q503X^ organoids at 2 months. Organoid #1 (n= 7511 cells), Organoid #2 (n= 9203), Organoid #3 (n= 9138). I. Graphical representation of downregulated GO-Terms in Patient^p.Q503X^ apical radial glia (aRG). Main biological process and cellular component GO-Terms are related to cell cycle and division. J. Graphical representation of upregulated GO-Terms in Patient^p.Q503X^ apical radial glia (aRG). Main biological process, cellular component and molecular function GO-Terms are related to neuronal differentiation and synapse formation. K. Violin plot of gene expression for CCPG1 (Cell Cycle Progression Gene 1) in each organoid for Patient^corrected^ and Patient^p.Q503X^.CCPG1 expression levels are higher in the aRG of Patient^p.Q503X^ organoids. **Violin plot of gene expression for MAP2 (Microtubule Associated Protein 2) in each organoid for Patient^corrected^ and Patient^p.Q503X^. MAP2 expression levels are higher in the aRG of Patient^p.Q503X^ organoids.**

To analyze this phenotype at higher resolution, we preformed single-cell RNA-sequencing analysis on a total of 50,592 cells from 6 individual organoids at 2 months (3 corrected and 3 haploinsufficient). To systematically perform cell type classification, we clustered cells from all organoids and compared signatures of differentially expressed genes with a pre-existing human fetal cortex single cell dataset^56^. This defined 9 main transcriptionally distinct cell types (Figure 4f; Suppl. Table 7), which included a large diversity of progenitors (apical radial glia, outer radial glia, and intermediate progenitor cells) and neuronal cell types (inhibitory neurons, corticofugal neurons and callosal projection neurons) representing all the main cell types of the endogenous human fetal cortex. Importantly we found that individual organoids were highly reproducibly in cell type composition consistent with previous reports^2,35^ (Figure 4g-h). This high level of reproducibility allowed us to preform differential gene expression analysis between Patient^Corrected^ and Patient^p.Q503X^ apical radial glia (table 8). In agreement with the observed increase in the differentiative division mode observed in haploinsufficient organoids we found an enrichment in terms related to neuronal differentiation, synapses and neuronal projection development. This was coupled with a downregulation of terms related to mitotic cell cycle process, cell division, mitotic spindle formation and centrosomes (Figure4i-l).

Overall, these data suggests that SYNGAP1 haploinsufficiency affect the balance of self-renewing versus differentiative division in apical radial glia cells.

### Decrease in SYNGAP1 levels leads to asynchronous cortical neurogenesis in vitro and in vivo

We next analyzed whether impairment in the division mode of hRGPs would affect the relative ratio of progenitor to neurons in 2-month-old SYNGAP1 haploinsufficient organoids. In line, with the observed acceleration in the timing of hRGCs differentiation, we found that the total percentage of SOX2+ cells were decreased in haploinsufficient organoids (Figure 5a, c; Suppl. Fig. 6a-b) and analysis of the percentage of NeuN+ cells showed an increase in the proportion of neurons generated (Figure 5b,d; Suppl. Fig. 6a,c). However, the percentage of TBR2+ cells remained consistent between genotypes (Suppl. Fig.6d-e). Interestingly, through single-cell sequencing analysis we found that the only cell population significantly altered was the apical radial glia which was decreased in patient organoids. Suggesting that at the 2-month old time point this might be the progenitor population most affected in Syngap1 haploinsufficient organoids (Table 9). However, the statistical comparison in the percentage of neurons in patient and corrected organoids was not significant. This discrepancy from single cell data and immunohistochemistry analysis might be due to the fact that single-cell sequencing may not accurately and reproducibly capture the increased percentage of neurons in patient organoids due to patient neurons having a more mature and therefore more fragile structure resulting in differential survivability after the dissociation step required in the single-cell pipeline.

**Figure 5.**
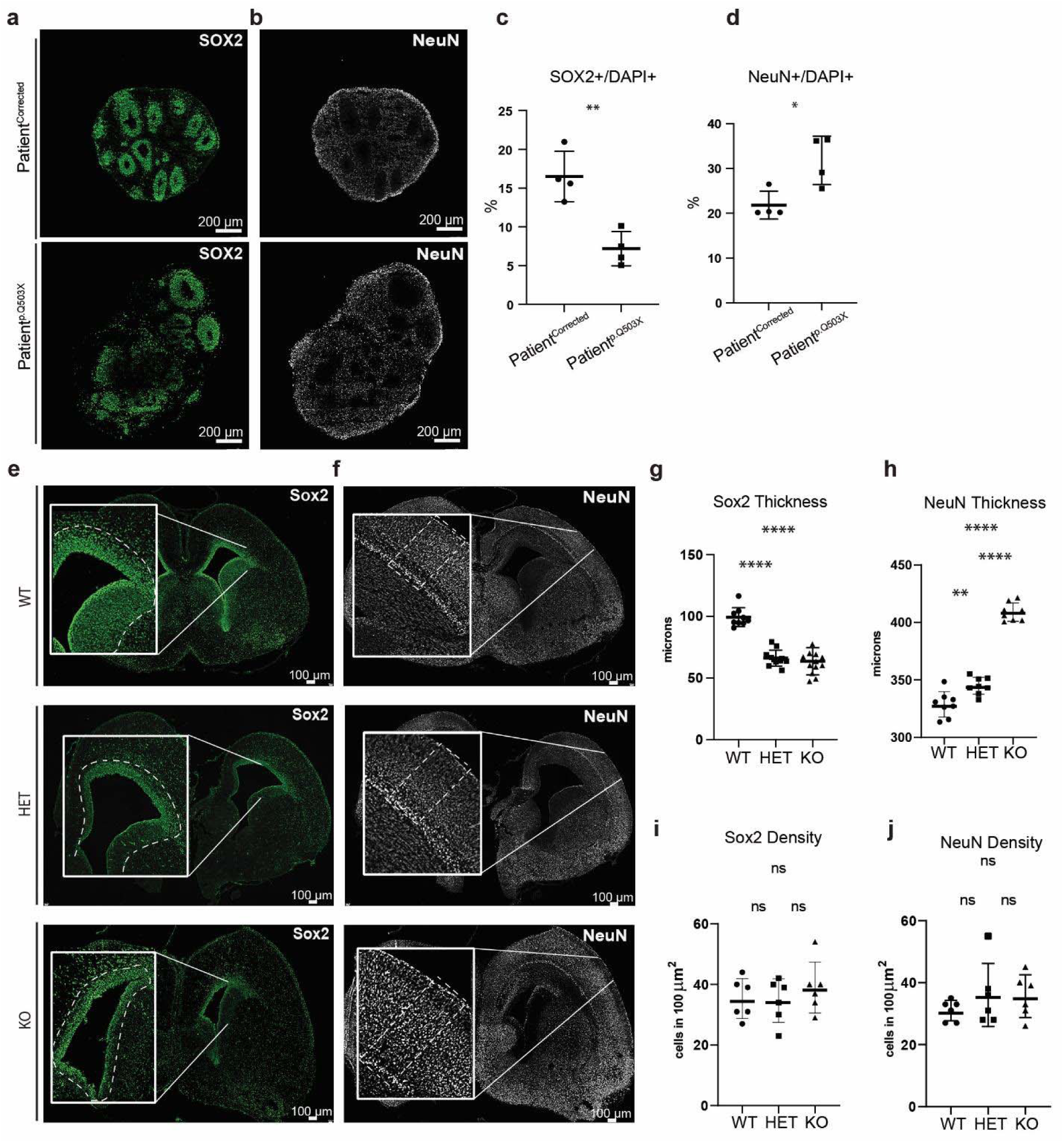
Decrease in SYNGAP1 levels lead to asynchronous cortical neurogenesis. A. Representative images of SOX2 expression in 2 month old Patient^Corrected^ and Patient^p.Q503X^ organoids. B. Representative images of NeuN expression in 2 month old Patient^Corrected^ and Patient^p.Q503X^ organoids. C. Quantification of the total number of SOX2 positive cells normalized to DAP1. Patient^p.Q503X^ organoids show a reduced density of SOX2 positive progenitors indicating and earlier depletion of the progenitor pool. Each dot represents an average value for all organoids from 1 differentiation. A Student’s t-test was performed on average values for 6-10 organoids from 4 differentiations. P Value =0.0032. Data is shown as mean ± SD. D. Quantification of the total number of NeuN positive cells normalized to DAP1. The higher density of NeuN positive cells in Patient^p.Q503X^ organoids indicates an accelerated differentiation of radial glial progenitors. Each dot represents an average value for all organoids from 1 differentiation. A Student’s t-test was performed on average values for 6-10 organoids from 4 differentiations. P Value =0.0184. Data is shown as mean ± SD. E. Representative images of SOX2 expression in SYNGAP1 Wild Type (WT), Heterozygous (Het) and KnockOut (KO) E18.5 mice. F. Quantification of SOX2 thickness lining the ventricular zone of the lateral ventricle in sections of E18.5 mouse brains. A decrease in the thickness of the SOX2+ area surrounding the ventricle in HET and KO as compared to WT mice indicates an earlier depletion of the proliferative niche. A One Way ANOVA was performed on 8-12 ventricle from four animals for each genotype. P<0.0001. G. Quantification of the number of SOX2^+^ cells in 100 um^2^ of the VZ area. The same density of SOX2+ cells was observed across genotypes. A One Way ANOVA was performed on 8-12 ventricles from four animals for each genotype. P=ns. H. Representative images of NEUN expression in SYNGAP1 Wild Type (WT), Heterozygous (Het) and Knock Out (KO) E18.5 mice. I. Quantification of NeuN thickness in the cortical plate in sections of E18.5 mouse brains. An increase in the NeuN thickness in KO and Het mice indicates an accelerated differentiation of radial glial cells. A One Way ANOVA was performed on 8 ventricles from four animals for each genotype. WT vs Het P=0.0311, WT vs KO P<0.0001, Het vs KO P<0.0001. J. Quantification of the number of NeuN^+^ cells in 100 um^2^ of the VZ area. A One Way ANOVA was performed on 8-12 ventricles from four animals for each genotype. P=ns.

As an additional line of evidence, we analyzed Syngap1 Het and KO mice at late embryonic stages (E18.5) to understand whether the developmental phenotype reported in organoids is conserved in vivo and across species. We stained sections from E18.5 mouse brains for the neural progenitor marker Sox2 and as found in organoids (Fig. 5e) we detected a reduction in the thickness of the size of the VZ in the Syngap1 Het and KO mice in comparison to WT littermates (Fig. 5g,i; Suppl. Fig. 6f-g). NeuN staining revealed a clear increase in the cortical thickness of the Syngap1 Het and KO mice illustrating an imbalance in the maintenance of radial glia and the production of cortical neurons compared to WT mice (Fig.5f,h,j). Interestingly, as observed in organoids, Tbr2 staining did not reveal any difference between genotypes (Suppl. Fig. 6h-j). Overall, the phenotypes observed in mice were in line with what was observed in organoids derived from patients with SYNGAP1 haploinsufficiency, indicating that a reduction in SynGAP1/SYNGAP1 protein levels impairs cortical neurogenesis in vivo as well as in vitro leading to the accelerated production of neurons. Interestingly, these changes in cell type composition did not significantly affect the overall organoid size over time (Suppl. Fig. S6k-l). This is consistent with the Patient^p.Q503X^ donor regularly scoring within the lower end of the normal head circumference size range according to the WHO child growth standard for males aged 0 to 2 years (Suppl. Fig. 6m). This lack of a clear microcephaly phenotype suggests that there are complex compensatory mechanisms at play in distinct progenitor populations at different developmental stages. Elucidation of this phenotype will require extensive characterization through lineage tracing approaches that can precisely reconstruct cell population dynamics and lineage relationships over time from individual cell types.

### SYNGAP1 haploinsufficient organoids exhibit accelerated maturation of cortical projection neurons

As additional evidence of the accelerated developmental trajectory in haploinsufficient organoids, we analyzed structural and functional features of neurons in intact 4-month-old organoids. Structural analysis of dendritic complexity in individual neurons showed a more developed dendritic tree arborization with an increase in the number of dendrites in haploinsufficient organoids (Fig. 6a-f). The higher degree of complexity in dendritic arborization was also supported by scRNA-seq data from 4-month-old Patient^Corrected^ and Patient^p.Q503X^ organoids (Suppl. Fig. 7c-e; Suppl. Table 10), showing in CFuPNs an upregulation of transcripts related to the functional categories of neuronal projection regulation and axon development including genes such as CNTN1, DCX, and SYT4 (Fig. 6g-h). CPNs showed enrichment in the transcription of genes including GRIN2B, DCX, and SLITRK5 with an overall upregulation of genes regulating neuronal projection morphogenesis and axon development (Fig. 6i-j). SLITRK5, a gene encoding for a protein known to suppress neurite outgrowth, was found to be downregulated, further supporting evidence of increased dendritic complexity. Importantly, upregulation in functional categories indicative of enhanced neuronal maturation was also observed in CFuPNs and CPNs within scRNA-seq data from 2-month-old Patient^Corrected^ and Patient^p.Q503X^ organoids, suggesting that this phenotype can be reproducibly detected across different batches and time points (Suppl. Fig. 7a-b; Suppl. Table 7). Functional neuronal maturation was monitored through recording of spontaneous neuronal activity using an Adeno-Associated Virus driving GCaMP6f as a proxy of intracellular calcium dynamics. Patient^Corrected^ and 03231^Control^ organoids displayed limited spontaneous basal calcium waves; however, haploinsufficient organoids showed an increase in network bursts, with faster transients, and coordinated bursts across distant GCaMP-positive soma (Fig.6k-n, Suppl. Fig.7f,h; Suppl. Video 3-6). Action potential (AP) generation and propagation were specifically blocked by tetrodotoxin (TTX), a voltage-gated sodium channel antagonist, and AP firing rate was found to be increased after bath application of glutamate, indicating that registered calcium waves are a result of glutamatergic synaptic connections (Suppl. Fig.7g,i). In line with previous evidence demonstrating precocious structural and functional maturation of SYNGAP1-deficient human glutamatergic neurons in 2D cultures^57^, these results indicate that haploinsufficient organoids exhibit higher levels of spontaneous activity and a more mature synchronized network.

**Figure 6.**
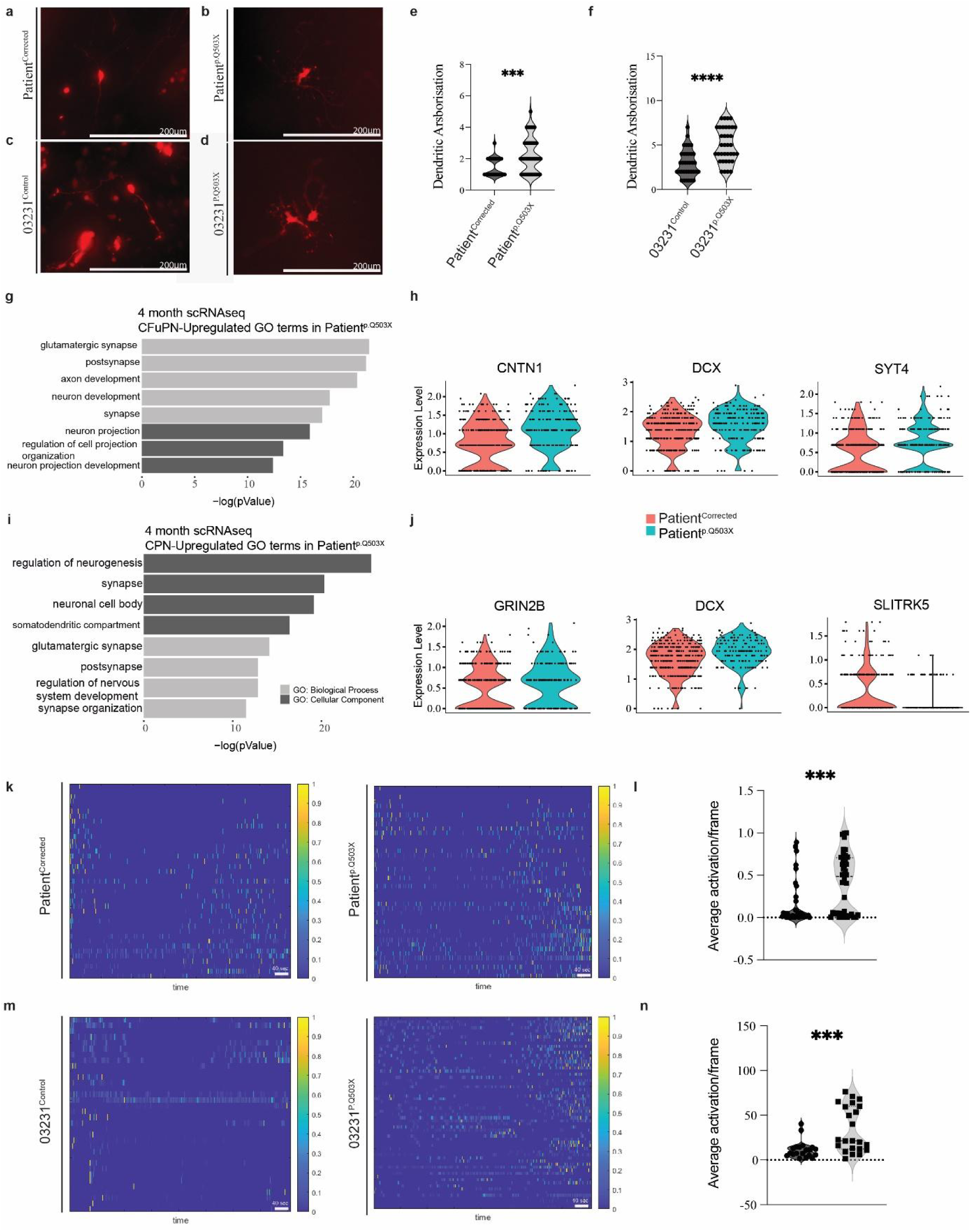
SYNGAP1 organoids exhibit accelerated maturation of cortical projection neurons. A. Representative image of RFP positive bipolar neuron from a 4-month-old Patient^Corrected^ organoid used for dendritic arborization analysis. B. Representative image of RFP positive multipolar neuron from a 4-month-old Patient^p.Q503X^ organoid used for dendritic arborization analysis. C. Representative image of RFP positive bipolar neuron from a 4-month-old 03231^Control^ organoid used for dendritic arborization analysis. D. Representative image of RFP positive multipolar neuron from a 4-month-old 03231^p.Q503X^ organoid used for dendritic arborization analysis. E. Dendritic arborization analysis on 4-month-old Patient^Corrected^ and Patient^p.Q503X^ organoids. Unpaired t-test performed on a total of 28 cells for Patient^Corrected^ organoids and 49 cells for Patient^p.Q503X^ organoids, from 3 independent experiments. P = 0.0009. F. Dendritic arborization analysis on 4-month-old 03231^Control^ and 03231^p.Q503X^ organoids. Unpaired t-test performed on a total of 35 cells for 03231^Control^ organoids and 36 cells for 03231^p.Q503X^ organoids, from 3 independent experiments. P = <0.0001. G. GO-Terms from single cell RNA sequencing preformed in 4 month old Patient^p.Q503X^ and Patient^Corrected^ organoids. Graphical representation of upregulated terms for Patient^p.Q503X^ Corticofugal Projection Neurons (CFuPN). Main biological process and cellular component GO-Terms are related to neuronal differentiation and synapse formation. H. Violin plot for selected genes showing expression levels between Patient^Corrected^ and Patient^p.Q503X^ organoids. DCX (Adj. P value = 0.04153089), CNTN1 (Adj. P value = 4.39E-07), SYT4 (Adj. P value = 6.77E-06) I. GO-Terms from single cell RNA sequencing preformed in 4 month old Patient^p.Q503X^ and Patient^Corrected^ organoids. Graphical representation of upregulated GO-Terms in Patient^p.Q503X^ Callosal Projection Neurons (CPN). Main biological process and cellular component GO-Terms are related to neuronal differentiation and synapse formation. J. Violin plot for selected genes showing expression levels between corrected and Patient^p.Q503X^ organoids. DCX (Adj. P value = 0.00040542), GRIN2B (P value =0.00031511), SLYTRK5 (P value = 3E-06). K. ΔF/F(t) from GCaMP6f2 recordings of Patient^Corrected^ and Patient^p.Q503X^ organoids visualized as a heatmap. L. Calcium average activation per frame analysis on 4-month-old Patient^Corrected^ and Patient^p.Q503X^ organoids, showing an increase in firing activity in Patient^p.Q503X^ organoids. Unpaired t-test performed on a total of 40 recordings in 12 Patient^Corrected^ and Patient^pQ503X^ organoids from 3 independent differentiations. P=0.0006. M. ΔF/F(t) from GCaMP6f2 recordings of 03231^Control^ and 03231^p.Q503X^ organoids visualized as a heatmap. N. Calcium average activation per frame analysis on 4-month-old 03231^Control^ and 03231^p.Q503X^ organoids, showing an increase in firing activity in 03231^p.Q503X^ organoids. Unpaired t-test performed on a total of 20 recordings in 6 03231^Control^ organoids and 24 recordings in 8 Patient^pQ503X^ organoids from 3 independent differentiations. P=0.0012.

Collectively, these data show that SYNGAP1 expression in hRGCs is required for precise control of their division mode, for the structural integrity and organization of the VZ, timing of neurogenesis, and proper cortical lamination. Finally, SYNGAP1 haploinsufficiency ultimately results in asynchronous cortical neurogenesis and accelerated maturation of cortical projection neurons.

## Discussion

Many human genetic studies of neurodevelopmental diseases including ASD have found enrichment of mutations in genes encoding classically defined synaptic proteins ^1,9,58-63^. Currently, most studies of SYNGAP1 and other synaptic proteins have been done within the context of mature synapses in rodent models^25^. Indeed, the lack of a reliable model to study the stage-specific functions of ASD risk genes during human brain development has limited our understanding of their role to rudimentary functional categories. Here, we leveraged 3D human brain organoids to perform longitudinal modeling and functional characterization of SYNGAP1, a top risk gene for ASD in the functional category of synaptic function. We detected SYNGAP1 in hRGCs and identified it as a key regulator of human cortical neurogenesis.

One advantage of using cultures of human cortical organoids is that, at early stages, they are composed by a relatively pure population of cortical progenitors. Analysis of the SYNGAP1 interactome within this population suggests an association between SYNGAP1 and the tight junction protein TJP1, which is a PDZ domain containing protein belonging to the MAGUK family. In neurons, SYNGAP1’s interaction with MAGUK proteins is key for protein localization within the PSD machinery^16-18^.Our results suggest a similar role of TJP1 in localizing SYNGAP1 function in the apical domain of radial glial progenitors. SYNGAP1 might regulate cytoskeletal organization in hRGCs through its RASGAP domain and its localization and assembly in macromolecular complexes through its PDZ ligand domain. This is particularly intriguing when considering the critical role of tight junctions and their association with the cytoskeleton in forming and preserving the apical junctional belt, which maintains radial glial apico-basal polarity and neuroepithelial cohesion^48^. As apical radial glial cells begin to differentiate, there is a downregulation of junctional proteins and a constriction of the apical junctional ring, which enables detachment from the ventricular wall and migration away from the VZ^64^. The disruption of junctional complexes regulating this delamination process has been shown to result in the collapse of apical RGCs morphology, disruption of the ventricular surface, and cortical lamination defects due to failed neuronal migration^65-68^. These phenotypes are consistent with several of our findings in SYNGAP1 haploinsufficient organoids, including disrupted ventricle formation and cortical lamination and the accelerated developmental trajectory of radial glial progenitors. Consistent with the high penetrance of pathogenic SYNGAP1 variants in patients, this phenotype was detected across two different genetic backgrounds and two different mutations.

Importantly, we also provide evidence for the expression and function of SynGAP1 in RGCs in vivo by exploring the earlier effects of the decrease in the level of SynGAP1 on the cortical progenitor dynamics in an embryonic mouse model of *Syngap1* mutation. Although murine neuronal progenitor biology differs from human^69^, we similarly observed an increased ratio of neurons to radial glial progenitors. Interestingly, decrease in SYNGAP1 levels have also been shown to regulate the ratios of neural progenitor cells to mature neurons in Xenopus tropicalis^21^, suggesting that Syngap1 controls the timing of cortical neurogenesis in vitro, in vivo, and across species.

As previously reported, developing neurons in SynGAP1 heterozygous mice exhibit accelerated structural and functional maturation, which have been linked to altered neural circuit connectivity and dysregulated network activity^40,70-72^. While it has been observed that SynGAP1 re-expression in adulthood can ameliorate certain relevant phenotypes induced by Syngap1 heterozygosity^72,73^, many core behavioral phenotypes in SynGAP1 haploinsufficient mice are not improved through adult-initiated gene restoration^71^. This phenomenon has been attributed to pleiotropic and temporally specific functions of the SYNGAP1/Syngap1 gene^73^. These current findings are consistent with this hypothesis because many of the Syngap1-related phenotypes described here are specific to early stages of brain development.

A growing body of literature suggests that a dysregulated neurogenesis program may lead to impaired neuronal wiring. This is because establishing proper neuronal circuits requires the spatiotemporally precise control of neuronal positioning, neurogenesis, and afferent and efferent synaptic connectivity^74,75^. Therefore, the impairment in neuronal excitability observed in SYNGAP1 patients may in part be explained by a disruption in microcircuit formation, driven by an altered developmental trajectory of cortical progenitors. Thus, our finding that cortical projection neurons in SYNGAP1 haploinsufficient organoids display accelerated development reshapes the current framework for therapeutic interventions, which could target not only the well-known alterations in synaptic transmission, but also the early developmental defects. The dual function of SYNGAP1 in progenitors and neuronal synapses underscores the importance of dissecting the role of ASD risk genes in specific cell types across developmental stages and suggests that a similarly nuanced approach may have broader relevance for studying other neurodevelopmental disorders (NDD). This is even more important considering that alterations in developmental trajectories during cortical neurogenesis are emerging as major contributors to the etiology of autism^2-8^.

In addition, human genetic studies have found enrichment of mutations in genes encoding PSD proteins^1,9,58,59,61-63^, which generally form large protein interaction networks that are considered risk factors for NDD^42,76,77^. In the synapses of the adult mouse brain, Syngap1 is a major hub in the PSD protein interaction network, and proteins associated with NDD are Syngap1 protein interactors^76,77^. We have found expression of multiple components of the core signaling machinery of the PSD in our proteomic analysis and shown that some of these proteins are interactors of SYNGAP1 in cortical progenitors. It is tempting to speculate that the complex controlling structural integrity at the PSD of excitatory neurons is also important in regulating the scaffolding properties of hRGCs. In support of this hypothesis, dysregulation of DLGAP4, another synapse-related scaffold protein, has been recently shown to regulate cell adhesion and actin cytoskeleton remodeling at the apical domain of radial glial cells during cortical neurogenesis in embryonic mice^78^. It will be crucial to characterize whether the disruption of proteins classically considered important for maintenance of the PSD scaffold in neurons may represent a point of convergence for NDD.

## Materials and Methods

### hiPSC Line Generation

#### Patient^p.Q503X^ Cell Line

An iPSCs line from a patient carrying a SYNGAP1 – c.1507C>T; p.Q503X nonsense mutation was generated. iPSCs were generated using the episomal expression of Yamanaka factors in patient-derived PBMCs^79^.

#### Patient^Corrected^ Cell Line

A sgRNA targeting the patient-specific mutation in SYNGAP1 was cloned into pSpCas9(BB)-2A-Puro (PX459) V2.0 (Addgene plasmid #62988). This, along with an HDR template containing the WT SYNGAP1 sequence, were nucleofected into the patient-derived iPSC line. Individual iPSC colonies were transferred to 24 well plates and subsequently underwent restriction enzyme-based genotyping. Positive colonies were then confirmed via Sanger sequencing and expanded in culture.

#### HDRTemplate

CCGCGAGAACACGCTTGCCACTAAAGCCATAGAAGAGTATATGAGACTGATTGGTCAGAAATATCTCAAGGATGCCATTGGTATGGCCCACACTCAGGCCCTCTTCTTCCCAAACCTGCCA.

The underlined CAG sequence corresponds to the insertion of the wildtype “T” base pair and the underlined T base corresponds to a silent mutation to disrupt the PAM sequence of the sgRNA. The substitution of the truncating “T” with the wildtype “C” base pair was screened for via restriction enzyme digestion and then confirmed via Sanger sequencing.

Mutation – c.1507C>T; p.Q503X nonsense mutation Genotyping info – Introduction of correction destroys DrdI restriction enzyme site which was used for initial screening of cell lines. Work on the patient-derived line patient p.Q503X is approved by the USC IRB: HS-18-00745.

#### RASGAP Dead Line

A sgRNA targeting the arginine finger region of SYNGAP1 was cloned into pSpCas9(BB)-2A-Puro (PX459) V2.0 (Addgene plasmid #62988). This, along with an HDR template to introduce the R485P RasGAP-dead mutation^80^, were nucleofected in the 03231 control iPSC line derived from a healthy 56-year-old male^45^. Individual iPSC colonies were transferred to 24 well plates and subsequently underwent restriction enzyme-based genotyping. Positive colonies were subsequently confirmed via Sanger sequencing. Guide Sequence used:

CGTGTTCTCGCGGAATATGHDR Template used: ACTTCCTTTCAGACATGGCCATGTCTGAGGTAGACCGGTTCATGGAACGGGAGCACtTaATATTCCcCGAGAACACGCTTGCCACTAAAGCCATAGAAGAGTATATGAGACTGATTGGTCAGA. The introduction of silent PAM mutation creates a new MseI restriction enzyme site which was used for initial screening of cell lines.

#### 03231^p.Q503X^

The 03231^p.Q503X^ cell line was generated in the 03231 control iPSC line derived from a healthy 56-year-old male ^45^. Via substitution of c.1507C>T; p.Q503X. A sgRNA targeting the SYNGAP1 c.1507C site was cloned into pSpCas9(BB)-2A-Puro (PX459) V2.0 (Addgene plasmid #62988). The sgRNA together with the HDR template to introduce the 1507C>T mutation. were nucleofected in the 03231 control iPSC line derived from a healthy 56-year-old male^45^. Individual iPSC colonies were transferred to 24 well plates and subsequently underwent restriction enzyme-based genotyping. Positive colonies were subsequently confirmed via Sanger sequencing. Guide Sequence used: CCATACCAATGGCATCCTTG.

#### HDR template used

CCGCGAGAACACGCTTGCCACTAAAGCCATAGAAGAGTATATG AGACTGATTGGT<u>TAG</u>AAATA<u>T</u>CTCAAGGATGCCATTGGTATGGCCCACACTCAGGCC CTCTTCTTCCCAAACCTGCCA

The Underlined TAG region shows the 1507C>T inserted mutation and the underlined T base shows the introduced silent PAM site.

Genotyping info – Introduction of correction modifies the DrdI restriction enzyme site which was used for the initial screening of cell lines.

Off-target predictions were carried out using CRISPOR^81^ he top ten off predicted off-target sites for the Patient 1-Corrected and RASGAP dead iPSC lines amenable to PCR were amplified using Q5 High-Fidelity DNA Polymerase (New England Biolabs; M0491S) and the resulting PCR products underwent Sanger Sequencing. All primers used for PCR amplification of predicted off-target sites were ordered from Integrated DNA Technologies and are listed in Supplemental Table 4.

### Cell Culture and Dorsal Forebrain Organoid Generation

hiPSC lines were maintained with daily media change in mTeSR (STEMCELL Technologies, #85850) on 1:100 geltrex (GIBCO, #A1413301) coated tissue culture plates (CELLTREAT, #229106) and passaged using ReLeSR (STEMCELL Technologies, #100-0484). Cells were maintained below passage 50 and periodically karyotyped via the G-banding Karyotype Service at Children’s Hospital Los Angeles. Organoid generation was performed as previously described in^35^.

### Procurement of Human Tissue

The de-identified human specimen was collected from autopsy, with previous patient consent to institutional ethical regulations of the University of California San Francisco Committee on Human Research. Collection was at a postmortem interval (PMI) of less than 24 hours. Tissue was collected at the following institution with previous patient consent to institutional ethical regulations: (1) The University of California, San Francisco (UCSF) Committee on Human Research. Protocols were approved by the Human Gamete, Embryo and Stem Cell Research Committee (Institutional Review Board GESCR# 10-02693) at UCSF. Specimens were evaluated by a neuropathologist as control samples. Tissues were cut coronally, and 1 mm tissue blocks were fixed with 4% paraformaldehyde for two days, and cryoprotected in a 30% sucrose gradient. The tissue was then frozen in OCT and blocks were cut at 30 μm with a cryostat and mounted onto glass slides.

### Procurement of Mouse Tissue

All animal procedures were conducted in accordance with the Guide for the Care and Use of Laboratory Animals of the National Institutes of Health, and all procedures were approved by the Institutional Animal Care and Use Committee (IACUC) of the Herbert Wertheim UF Scripps Institute for Biomedical Innovation and. Males and females were used in all experiments and final male/female ratio in datasets reflect uncontrollable variables, such as the ratio of male/female (M/F) offspring achieved from the multigenerational breeding schemes and experimental attrition. Mice were housed four or five per cage on a 12-h normal light–dark cycle.

We used inbred *Syngap1* constitutive *Syngap1*^+/−^ (heterozygous) mice^13^. Each line is maintained by colony inbreeding on a mixed background of C57-BL6/129s. Every seventh generation, *Syngap1*^+/−^ mice are refreshed by crossing colony breeders into C57-BL6/129 F1 (Taconic B6129F1) animals for one generation. Offspring from these crosses, like those used for this study, are then inbred for up to seven generations. For staging of embryos, the day of vaginal plug was considered E0.5. Embryo collection was carried out at E18.5. The dams were anesthetized using an isoflurane induction chamber (5% isoflurane) and placed in a nose cone with flowing isoflurane for maintenance (2.5% isoflurane). After fixation on a prewarmed surgery platform and sterilization of the skin with Betadine, the abdominal cavity was opened through 2 incisions along the midline to expose the uterine horns. The uterine wall was incised to expose embryos within their fetal membranes. The pups were extracted from the yolk sac and then placed on a petri dish with ice cold PBS. All pups were decapitated, the brains quickly extracted from the skull, and each submerged in vials with 4% PFA. Following the pup dissections, dams were euthanized by cervical dislocation while under isoflurane anesthesia. Genotyping of all transgenic mouse lines was outsourced to Transnetyx automated genotyping services.

### Singular Neural Rosette Tissues

All pluripotent lines were maintained in Essential Eight medium (E8) (Thermo Fisher, #A1517001) on geltrex (GIBCO, #A1413301) coated tissue culture plates and routinely passaged with ReLeSR (STEMCELL Technologies, #05872). RGCs derivation from hPSCs was performed using Essential Six Medium (E6) (Thermo Fisher, #A1516501)^82^. To generate RGCs-derived micropatterned, cells were first rinsed with PBS, dissociated with Accutase (STEMCELL Technologies, #07922) for 5 min at 37°C, and collected via centrifugation at 1000 rpm for 5 min. Singularized RGCs were re-suspended in E6 media with 10 μM ROCK inhibitor (Y27632; STEMCELL Technologies, #72302) and seeded onto micropatterned substrates at 75,000 cells/cm2 in 2 mL of media per well. The following day, the media was replaced with 2 mL of E6 media, and 50% media changes were performed daily thereafter. 96-well plates were custom made with micropatterning of poly(ethylene glycol methyl ether)-grafted substrates, presenting arrays of 250μm diameter circular regions and coated with Matrigel over-night^44^.

### Immunohistochemistry

#### Organoids

Organoids were fixed in 4% PFA for 30 min at room temperature before an overnight incubation at 4L in 30% sucrose solution. Organoids were then embedded in Tissue-Tek O.C.T. compound (Sakura, #62550) and sectioned at 20 µm with a cryostat onto glass slides (Globe Scientific, #1354W). Slides were washed 3x with a 0.1% Tween20 (Sigma, #P9416) solution before a 1-hour incubation in 0.3%TritonX-100 (Sigma, #T9284) and 6% bovine serum albumin (Sigma, #AA0281) solution. An overnight incubation at 4L in a primary antibody solution was followed by a 2-hour room temperature incubation in a secondary antibody solution, both consisting of 0.1%TritonX-100 and 2.5%BSA with 3 washes before and after secondary antibody incubation. Slides were cover slipped using Fluoromount G (EMS, #50-259-73). BrdU staining was performed by initially treating the slides with 2M HCL (Sigma, H1758-100ml) for 30 minutes at room temperature followed by a neutralization step with 0.1 M sodium borate (Millipore, # SX0355-1) buffer at pH 8.5 for 10 minutes at room temperature before continuing with the above described IHC protocol.

#### Mouse Tissue

Mouse tissue underwent a similar IHC protocol aside from the addition of antigen retrieval prior to beginning the IHC protocol. Antigen retrieval was performed by incubating slides in citrate buffer at 95L for 30 minutes. Slides were then returned to room temperature and allowed to sit in the citrate buffer for 1 hour before continuing with the IHC protocol.

| **Antibody** | **Species** | **Catalog Number** | **Company** |
| --- | --- | --- | --- |
| DAPI | -- | 80051-386 | Sigma/Millipore |
| SATB2 | Mouse | AB51502 | Abcam |
| MAP2 | Chicken | AB5392 | Abcam |
| NESTIN | Mouse | AB22035 | Abcam |
| CTIP2 (BLC11B) | Rat | AB18465 | Abcam |
| SOX2 | Goat | AF2018 | RD Systems |
| TBR2 (EOMES) | Rabbit | AB2283 | Millipore |
| TBR2 (EOMES) | Chicken | AB15894 | Millipore |
| PAX6 | Rabbit | 901301 | Biolegend |
| Pericentrin | Rabbit | ab4448 | Abcam |
| Anti-RFP | Rabbit | 600-401-379 | Rockland |
| p-Vimentin | Mouse | D076-3 | MBL |
| ZO-1 (TJP1) | Mouse | 610966 | BD Transduction Laboratories |
| SYNGAP1 | Rabbit | 5539 | Cell Signaling Technology |
| SYNGAP1 | Rabbit | 5540 | Cell Signaling Technology |
| SYNGAP1 | Rabbit | Ab77235 | Abcam |
| Acetylated Tubulin | Mouse | T6793-100UL | Sigma Aldrich |
| Bromodeoxyuridine (BrdU) | Rat | AB6326 | Abcam |
| NeuN | Rabbit | AB177487 | Abcam |

### Peptide competition assay

Three consecutive organoids slices were used for analysis. Rabbit anti-SYNGAP1 (5539, Cell Signaling Technology) was diluted at a concentration of 1:200 in 0.3%TritonX-100 (Sigma, #T9284) and 6% bovine serum albumin (Sigma, #AA0281) blocking solution. The solution was divided in three tubes where no blocking antigenic peptide, 5x and 10x peptide (32835S, Cell Signaling Technology) were added. Tubes were incubated with agitation overnight at 4L. The immunofluorescence procedure was performed as described in the manuscript.

### BrdU Labeling in Cortical Organoids

2-month-old organoids were incubated with 10 µm of the thymidine analogue Bromodeoxyuridine (BrdU) (Sigma, #B5002) for 2 hours before being rinsed with PBS and then allowed to remain in culture for 24hrs. After the 24hr chase organoids were removed from culture and fixed for 30 minutes in 4% PFA at room temperature.

### Whole proteome analysis of organoids

Deep proteome profiling was performed on 21 human cortical organoids at D.I.V. 7 from the <u>03231</u> control line. After lysis in RIPA buffer (Pierce), extracted proteins were reduced with 5 mM DTT, followed by incubation with 20 mM iodoacetamide in the dark. Sample cleanup was performed using the SP3 method ^83^ and on-bead digestion was performed overnight using Trypsin/Lys-C (Promega). Eluted tryptic peptides from SP3 beads were pre-fractionated by high pH reverse phase chromatography into 96 individual fractions using a 64-minute gradient from 1% to 50% solvent B (A: 10mM NH_4_OH, B: acetonitrile) on a Cadenza C18 column (Imtakt) The collected 96 fractions were recombined into final 24 fractions by pooling every 24^th^ fraction for LC-MS/MS analysis. 500 ng peptide from each fraction was analyzed by a 60-minute LC/MS/MS method on an Orbitrap Fusion Lumos mass spectrometer (ThermoFisher Scientific) interfaced with a Ultimate 3000 UHPLC system (ThermoFisher Scientific). Full scans were acquired in the Orbitrap at a resolution of 120K and a scan range of 400-1600 m/z. Most abundant precursor ions from the full scan were selected using an isolation window of 1.6 Da and subjected to HCD fragmentation with an NCE of 30% and detection in the iontrap. Raw data files were searched using Byonic (v2.16.11) in Proteome Discoverer (v2.4) against the Swissprot human protein database (downloaded November, 2020). The following search parameters were used: fully tryptic peptides with a maximum of 2 missed cleavages, 10 ppm precursor mass tolerance, 0.5 Da fragment mass tolerance, fixed modification of cysteine carbamidomethylation, oxidation of methionine, deamidation of glutamine and asparagine were selected as dynamic modifications. The protein and peptide-level confidence thresholds were set at 99% (FDR <0.01). Using this pipeline, we identified a total of 8686 proteins, including 24 unique peptides for *SYNGAP1*. The protein/gene list obtained from MS analysis was imported into SynGO portal and cellular component annotation where obtained.

### Generation and Analysis of the SYNGAP1 Interactome

SYNGAP1 protein immunoisolation was performed using 50 human cortical organoids at D.I.V. 7, generated from both the 0323^control^ and the Patient^Corrected^ iPSC lines, in duplicate assays. SYNGAP1 was immunoprecipitated using 2ug/ml of SYNGAP1 antibody (Cell Signaling 5540) and 3mg/ml of total protein lysate. The SYNGAP1 KO cell line was used in duplicate assays a IP control. Peptide identification was performed as described for whole proteome analysis. Protein interactors were defined as proteins present in duplicate samples and absent in KO controls.

### Quantitation of SYNGAP1 total protein and identification of the SYNGAP1 alpha 1 isoform

Peptides were synthesized commercially (ThermoFisher Scientific) TVSVPVEGR; DAIGEFIR, and GSFPPWVQQTR. SYNGAP1 id: A0A2R8Y6T2. Both unlabeled light and isotope labeled heavy forms were synthesized. Commercially synthesized labeled peptides used ^13^C and ^15^N labeled lysine or arginine. An internal standard was prepared from the isotope-labeled standards in 3% acetonitrile with 0.1% formic acid with 5 fmol/µL of Peptide Retention Time Calibration (PRTC) mixture (ThermoFisher Scientific) and 10 µg/mL E. coli lysate digest (Waters) as a carrier. Isotope-labeled peptides were dissolved at 2000 pg/mL. Standard Concentration: 1000pg/mL. A calibration curve was prepared from the light standards in 3% acetonitrile with 0.1% formic acid and 10 µg/mL E. coli lysate digest, but without the PRTC mixture. The highest concentration stock was prepared at 250000 pg/mL The final calibration standards were made by mixing the stocks 1:1 with internal standard mixture. Dried protein digests were dissolved in 20µL of a 1:1 mixture of internal standard mixture and 10 µg/mL E. coli lysate digest and 5 µL was injected on-column for PRM data acquisition.

Samples were analyzed on an Ultimate 3000 nanoflow UHPLC system coupled to an Orbitrap Fusion Lumos mass spectrometer (ThermoFisher Scientific). Digested peptides were separated using a 25 cm C18 EasySpray column (75 µM ID, 2 µm particle size) using 0.1% formic acid in water as mobile phase A and 0.1% formic acid in acetonitrile as mobile phase B. Peptides were analyzed using a timed parallel reaction monitoring (tPRM) method. Expected retention times were measured before each batch by analyzing the internal standard. Each peptide was given a retention time window of ±2 minutes. Precursor m/z for each peptide. Data was analyzed in Skyline. For each assay, top three fragment ions without any interference were selected for quantitation. The calibration curve used the ratio of light to heavy peptide using a bilinear curve fitting and 1/x^2^ weighting. The limit of quantitation was estimated as the lowest calibration point with a coefficient of variability below 15% and an average error below 15%. Limit of detection was calculated using the standard of deviation of the blank and the standard of deviation of the sample at the limit of quantitation; the higher value was used as the limit of detection. Peptides were considered detected if all three product ions were detected, the dot product ratio was at least 0.7, and the measured quantity was above the limit of detection.

### Bulk-RNA Sequencing

Organoids derived from haploinsufficient and corrected control lines were collected at DIV 7 from 2 independent differentiations (50 corrected and 100 patient organoids per genotype, per differentiation). Total RNA was isolated using Qiagen columns. Library preparation and RNA sequencing were performed as a service by QuickBiology.

Data preprocessing was performed with *trimmomatic* to filter out adapter sequences and low-quality reads. The processed reads were mapped to the human reference genome from Ensembl (GRCh38.p13) using *<u>HISAT2 v2.1.0</u>*. We summed the read counts and TPM of all alternative splicing transcripts of a gene to obtain gene expression levels and restricted our analysis to 20000 expressed genes with an average TPM >1 in each sample. Differential expression analysis was performed with the *<u>DESeq2 package</u>* (v1.20.0). The following cutoffs values were used for assigning differentially expressed genes (DEGs): P-adjusted valueL<L0.05, false discovery rate (FDR) < 0.05 and |log2FC| ≥ 0.6. We obtained a list of both upregulated and downregulated DEGs between SYNGAP1+/-and control organoids. ClusterProfiler software (v.4.2.2) was used to perform functional annotations of the DEGs, according to <u>Gene Ontology</u> (GO) categories (biological process, molecular function and cellular components). Using these gene lists, we searched the Panther GO-Slim Biological Processes ontology database using a statistical overrepresentation test (FDR, P < 0.05).

### Single Cell Dissociation

7-day-old organoids were rinsed with 1X PBS (Corning, #21-040-CV) and incubated at 37L for 15 min with 1x TripLE Express Enzyme (Thermo Fisher Scientific, #12-605-010). Organoids were dissociated by pipetting up and down with a 1000 µl pipette followed by 200 µl pipette until a single cell resuspension was obtained. Cells were plated onto 1:100 geltrex coated round plastic coverslips (Thermo Fisher Scientific, #NC0706236) at a density of 15800 cells/cm^2^.

### Single Cell Dissociation for scRNA-Sequencing

Organoids were dissociated as previously described^30,51^. Briefly, 3-4 pooled organoids at 3-4 months of age or three single organoids at 2-month age from both haploinsufficient and corrected lines were transferred to a 60 mm dish containing a papain and DNase solution from the Papain Dissociation Kit (Worthington, #LK003150). Organoids were minced into small bits with razors and incubated for 30 min at 37°C on an orbital shaker, then mixed with a 1ml tip several times and returned for another 5-10 min at 37°C. Next, the pieces were triturated 5-10 times with a 10 ml pipet and transferred to a 15 ml conical tube containing 8 ml final Inhibitor solution and DNase. The tube was inverted a few times and left upright for a few minutes for larger debris (if any) to settle, then the single cell suspension was transferred to a new conical tube, and centrifuge for 7 min at 300 g. The cell pellet was resuspended in 500 ul to 1 ml of 0.04% BSA/PBS and passed through a 0.4 μm filter basket and counted. The solution was then adjusted to a target concentration of 1000 cells/µl.

### Sequencing Analysis, Quality Control (QC), and Clustering

10X Genomic scRNA sequencing was performed by the USC Molecular Genomics Core Facility. Samples were processed on the 10x Single Cell Gene Expression 3’ v3.1 kit. Raw sequencing reads were aligned with the human reference genome (GRCh38-2020-A) via the CellRanger (v6.0.2) pipeline to generate a cell-by-gene count matrix. Next, we used Seurat (v4.0.1 using R v4.0.) to perform initial QC with standard cutoffs of min.cells = 3, min.features = 200, mitochondrial percentage (<15%), and removal of low complexity cells (nCount < 1250). For the 2-month age organoids, we recovered, after quality control, a total of 25832 patient and 24740 corrected cells (total: 50572), while for the 4-month, pooled organoids, 3123 patient and 7540 corrected cells (total: 10663) were recovered. The regression of the cycling cell genes, normalization, variable features, and scaling were done using the SCTransform function. Principal component analysis (PCA) was performed. No batch correction was required for the merging of the six individual organoids from patient and correct lines. However, comparison of our organoids with other human fetal and organoids data at 2-month and 4-month^84^ required merging using the Seurat pipeline based on Canonical Correlation Analysis (CCA) integration. Next, using the FindNeighbors and FindClusters functions (resolution = 0.2, 0.8 and 1.2), clusters of cell types were generated. Clusters were classified according to known markers, previously identified molecular profiles from organoids^34,35^, a human fetal data set^56^ and coclustering (CCA) with other 2-month and 4-month old organoids^84^. To calculate the DEGs between control and mutant organoids for each cluster, we used the *FindMarker* function with the test.use attribute set to the default Wilcoxon rank sum test. Using the ToppFun application on the Toppgene site (toppgene.cchmc.org), we entered the significant DEGs from each cell type along with their respective background genes to detect and generate GO-terms with functional enrichment. Next, we selected representative genes from relevant and significant GO terms and visualized them as violin plots with an equal sample size between conditions.

### RFP Labeling

Organoids were transduced with EF1A-RFP lentivirus (Cellomics Technology, # PLV-10072-50). 1 µl of stock virus (1×10^8 TU/ml) was diluted into 500 µl Cortical Differentiation Medium IV (CDMIV, without Matrigel) in a 24-well ultra-low attachment plate (Corning, # 3473) containing a single organoid. After 24 hours of incubation, full media change was performed. 48 hours later, organoids were transferred to a 6-well ultra-low attachment plate (Corning, #3471). 1 week after transduction, organoids were randomly selected for imaging analysis and individually transferred to a u-Slide 8-well Glass-bottom plate (Ibidi, #80827).

### Calcium Imaging

Organoids were transduced with pAAV-CAG-SomaGCaMP6f2 (Addgene, #158757) as described in^2^. Four-month-old cortical organoids were randomly selected and transferred to a recording chamber kept at 37L°C using a heating platform and a controller (TC-324C, Warner Instruments) in 5% methyl-cellulose in BrainPhys Imaging Optimized Medium (STEMCELL Technologies, #05796). Imaging was performed using a SP-8X microscope with a multiphoton laser. Time-lapse images were acquired at 1 frame for 860 ms, using a 25x 0.95 NA water objective (2.5 mm WD) and resulting in a view of 200 x 200 µm^2^. Basal activity was recorded for 10 mins in each of the 3 randomly selected areas of the imaged organoid. Pharmacological treatment was performed with a bath application of Tetrodotoxin, TTX (Tocris, #1078/1) at a final concentration of 2 µM, and glutamate (Hello Bio, #HB0383) at 100 µM.

### Singular Neural Rosette Tissues Analysis

The presence of 0 or more than 1 rosette was assessed and classified as 0 rosettes.

### Singular Neural Rosette Live Imaging Analysis

Singular rosettes have been imaged with a 20x objective of Leica Thunder Microscope for 14 hours under brightfield light. Co2 and temperature were maintained at 5% and 37C throughout the recording using a recording chamber.

### Morphological Dendrite Analysis

RFP positive organoids were imaged using a 20x objective with the Leica Thunder Microscope. Maximum projection of each organoid was applied, and the number of dendrites per cell was calculated using the Sholl Analysis plugin in Image-J software.

### Ventricular Zone Analysis of 2-month-old Organoids

Two-month-old organoids were sectioned and immunostained for SOX2 and DAPI. Only cryosections near the middle of each organoid were used for analysis. The ventricular zone (VZ) was defined by exclusive SOX2 immunoreactivity and neural tube-like morphology. Image-J software was used to analyze the thickness, area, and total number of VZs. The line tool was used to measure the thickness of each VZ (µm); an average of 3 measurements per VZ were considered. The freehand tool was used to trace the entire VZ and measure the total area (µm^2^). For each organoid, 3-6 regions of interest (ROI) were defined, and the number of VZs in each Region of Interest (ROI) were counted. For all VZ analyses, 4 independent differentiations and 3 organoids from each differentiation were measured.

The organized versus disorganized analysis of the VZ was based on MAP2 staining. The ventricles with clusters of MAP2 positive cells in the germinal zone were defined as disorganized; the ventricles showing clear boundaries between the cortical and germinal zones were defined as organized. For each line, 3 organoids from 3 independent differentiations were quantified.

The cleavage angle was defined as the angle between the ventricular apical surface and the cleavage plane of dividing cells. Pericentrin and pVimentin were used to stain the centrosome and the dividing RG, respectively, to clearly visualize the cleavage plane. The angle was measured manually with the angle tool in Image-J.

Binning analysis was performed on relatively isolated ventricles to ensure the binning area has no other interfering ventricles. Seven 25×50 µm rectangular bins were stacked vertically starting from the edge of the ventricle and extending to the nearest edge of the organoid. This provided a uniform grid or binning area that was then used as a visual for the layering of specific cell markers including: CTIP2, SATB2, and TBR2. Positive cells for each stain in each of the bins were manually counted and normalized to the total number of DAPI positive cells per bin.

### NeuN, TBR2 and SOX2 Density Analysis of 2-month-old Organoids

2-month-old organoids stained by anti-NeuN, TBR2, and SOX2 were imaged and used for quantification. Images were opened in ImageJ software and the background noise was reduced. To count all the cells, DAPI-positive cells were initially counted. The lowest and maximum Threshold values were set at 30 and 250, respectively. The proportion of NeuN, TBR2 or SOX2-positive cells was calculated by normalization to the number of DAPI-positive cells.

### NeuN, TBR2 and SOX2 Thickness and Density Analysis of E18.5 Mouse Tissue

For each genotype (WT, HET, KO), the thickness of labeled cells within the lateral cortex was quantified in ImageJ software from 4-5 20μm sections sampled at ∼100μm intervals along the rostro caudal axis within the presumptive somatosensory region of the cortex.

Density was calculated by the number of NeuN, TBR2, or SOX2 positive cells in 100 um^2^.

### SOX2 Area and Density Analysis of E18.5 Mouse Tissue

For each genotype (WT, HET, KO), the SOX2 positive area around the VZ was selected by free-hang image tool in ImageJ software and quantified from 4-5 20μm sections sampled at ∼100μm intervals along the rostro caudal axis within the presumptive somatosensory region of the cortex.

### BrdU Pulse-Chase Data Analysis

Binning analysis was performed on relatively isolated ventricles to ensure the binning area has no other interfering ventricles. Seven 25×50 µm rectangular bins were stacked vertically starting from the edge of the ventricle and extending to the nearest edge of the organoid. This provided a uniform grid or binning area that was then used as a visual for the layering of specific cell markers including: SOX2, NeuN, and BrdU. Positive cells for each stain in each of the bins were manually counted and normalized to the total number of DAPI positive cells.

#### Organoid size

Every three days, bright-field microscopy was utilized to capture images of all organoids from day 3 to 60. ImageJ software was then used to measure the area and the perimeter of each single organoid. The Prism software was then used to plot the size average of the organoids of each differentiation.

#### Organoid yields

To check the organoids’ survivability, bright-field microscopy was used to image the organoids during the culture period. All organoids were then counted. Measurement of survival in percentile was made considering the number of starting organoids.

### Calcium Imaging Data Analysis

Raw tiff calcium imaging files were analyzed using the CNMF and CalmAN package as previously described (CaImAn an open-source tool for scalable calcium imaging data analysis^85^, to identify fluorescent transients and spike estimation in MATLAB (MathWorks). Calcium traces were plotted as relative scaled height in function of time.

### Statistical Analysis

Data is shown as mean ± SD/SEM. Statistical analysis was performed using the Graph Pad Prism

6.0. Shapiro-Wilk test was performed to determine the normality of the data (alpha=0.05). Comparisons of means in 2 or more groups were made using an unpaired Student’s t-test, analysis of variance (ANOVA) or Chi-square test. Significant main effects were analyzed further by post hoc comparisons of means using Bonferroni’s multiple comparisons test.

### Ethical approval

Work on the patient-derived line patient p.Q503X is approved by the USC IRB: HS-18-00745

## Acknowledgments

We thank the Syngap Research Fund and families participating in this study for their collaboration. We thank the Pediatric Neuropathology Research Lab (PNRL) of UCSF and honor the families who generously donated the tissue samples used in this study.

We thank Dr. Paola Arlotta, Dr. Fiona Francis, current and former members of the Quadrato lab for insightful discussions and feedback on this project. We thank Cristy Lytal for editing the manuscript.

We thank Seth Walter Ruffins and the Optical Imaging Facility at USC for providing guidance and support of imaging analyses. We thank Christopher Taitano-Johnson, Thomas Rintoul and Angela Albanese for outstanding technical support.

## Author contributions

M.B., A.D., T.N., M.P.C., and G.Q. conceived the experiments. M.B and A.D. generated, cultured, and characterized single rosette cultures; J.P.U. performed live imaging of single rosettes; G.K. and R.S.A. prepared single rosette culture substrates; M.B., A.D.,T.X, N.H. and S.N. generated, cultured, and characterized all the organoids used in this study. A.D. and M.B. performed scRNA-seq and bulk RNA-seq experiments with help from T.X., A.A. and T.N.; T.N. performed scRNA-seq analysis and worked on cell type assignments and data analysis; T.X. performed bulk scRNA-seq analysis. B.W. performed sample preparation for Proteomic analysis, and R.M., R.S., P.P. performed analysis of the proteomic data under the supervision of M.P.C.; M.B. performed the calcium imaging experiments and analysis; B.W. designed and generated the SYNGAP1 p.Q503X, corrected, RGD, and 03231/SYNGAP1 P.Q503X-edited lines under the supervision of M.P.C.. E.H. prepared and provided human fetal tissue. C.R. prepared and provided mouse fetal tissue under the supervision of G.R. G.R. contributed to the data interpretation. G.Q. supervised all aspects of the project; M.B., A.D., and G.Q. wrote the manuscript with contributions from all authors.

## Data and materials availability

Bulk RNA Seq Raw Counts https://figshare.com/s/fe1ab60f2a3ebcaae7ff

2-month-old scRNA-seq data

Patient^Corrected^ organoid 1 https://figshare.com/s/93d4d3b71ec8d9951eb4

Patient^Corrected^ organoid 2 https://figshare.com/s/7603008aefc936fd53d7

Patient^Corrected^ organoid 3 https://figshare.com/s/d19a8905e1296e2055bd

SYNGAP1^p.Q503X^ organoid 1 https://figshare.com/s/794cfe68f8caedfbd440

SYNGAP1^p.Q503X^ organoid 2 https://figshare.com/s/d5a9e7d3ab45b3c50a21

SYNGAP1^p.Q503X^ organoid 3 https://figshare.com/s/05dc105003079568bfc3

4-month-old scRNA Seq data

SYNGAP1^p.Q503X^ organoids matrix.mtx.gz https://figshare.com/s/1cd0dc51bcf6f1028cd2

SYNGAP1^p.Q503X^ organoids features.tsv.gz https://figshare.com/s/c08cfabef967271dcfbb

SYNGAP1^p.Q503X^ organoids barcodes.tsv.gz https://figshare.com/s/2b493cc0108d5f2cbc35

Patient^Corrected^ organoids matrix.mtx.gz https://figshare.com/s/5b8b4b0cd4818c9c2e7d

Patient^Corrected^ organoids features.tsv.gz https://figshare.com/s/d6054204bd0514a87b5e

Patient^Corrected^ organoids barcodes.tsv.gz https://figshare.com/s/c1684604c58b0b04ba05

Proteomics IDEntification Database (PRIDE). Username. Password: oWG3mX6A. Project accession: PXD034090.

---

**Supplementary Figure 1.**
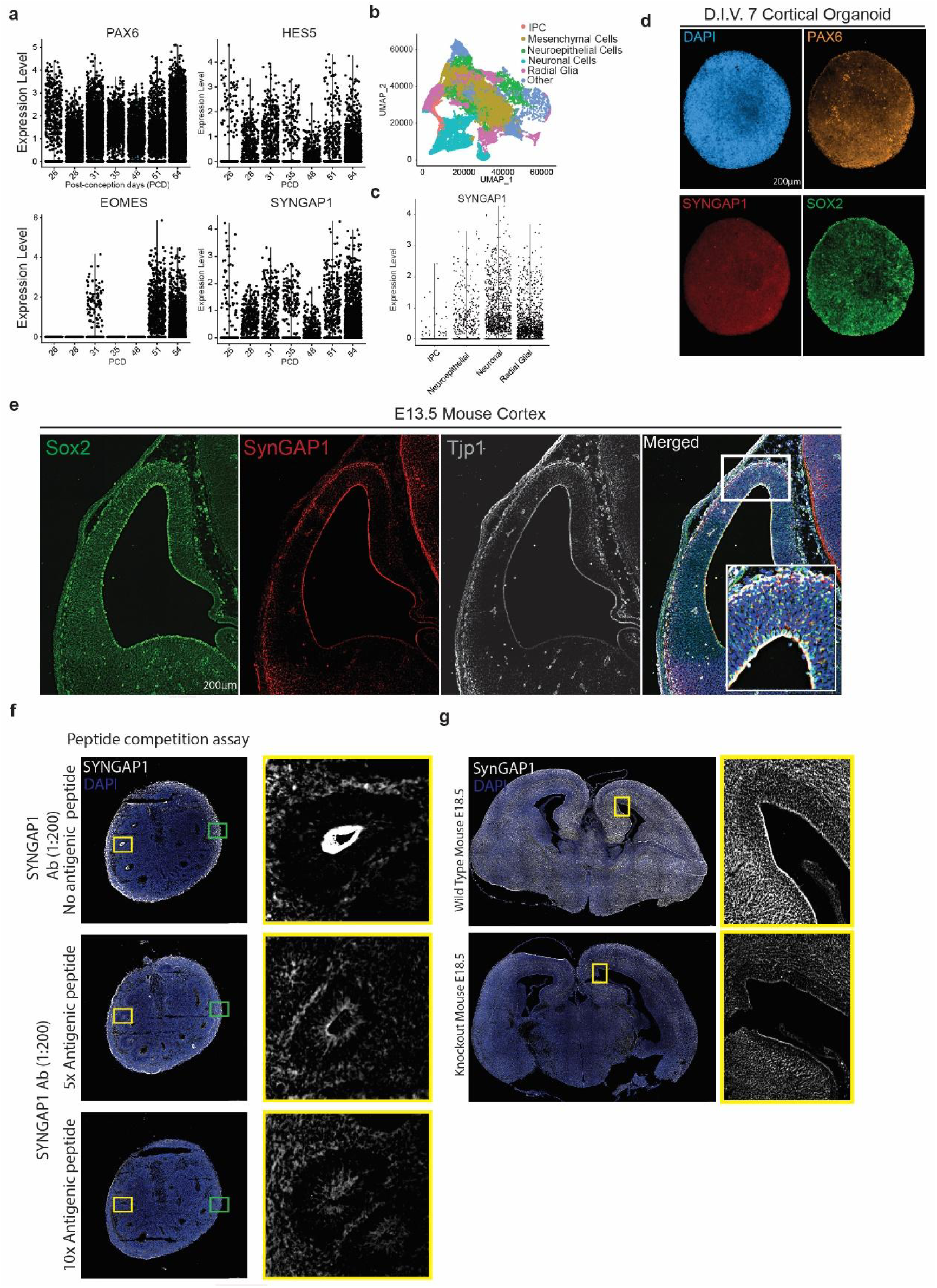
A. Expression of early forebrain marker genes of PAX6, HES5, EOMES (TBR2) and SYNGAP1 from post-conception day (PCD) 26 to 54 from single cell RNA-seq data. B. UMAP visualization of age-dependent clustering of fetal single cells. C. SYNGAP1 expression at PCD 56 grouped by cell types; intermediate progenitor cells (IPC), neuroepithelial cells (NE), radial glial cells (RGCs) and neurons. D. D.I.V. 7 cortical organoids are composed of cells positive for the neural stem cell marker SOX2, the radial glial progenitor marker PAX6, the nuclear marker DAPI and SYNGAP1. E. A coronal section from E13.5 mouse brain showing expression of the neural stem cell marker SOX2, the tight junction protein TJP1, and SYNGAP1. SYNGAP1 is highly expressed at the ventricular wall. White box indicates the Region of Interest selected for the merged images showing colocalization of DAPI, TJP1, and SYNGAP1. F. Peptide competition assay shows the specificity of the SYNGAP1 antibody used. 5X and 10X concentrations of the commercial antigenic peptide were evaluated, showing a strong reduction in specific signal in the apical wall of the ventricular zone. G. SynGAP1 expression in E18.5 wild type and SynGAP1 KO mouse showing the overall decrease in SynGAP1 levels. Decreased levels of SynGAP are most evident at the VZ.

**Supplementary Figure 2.**
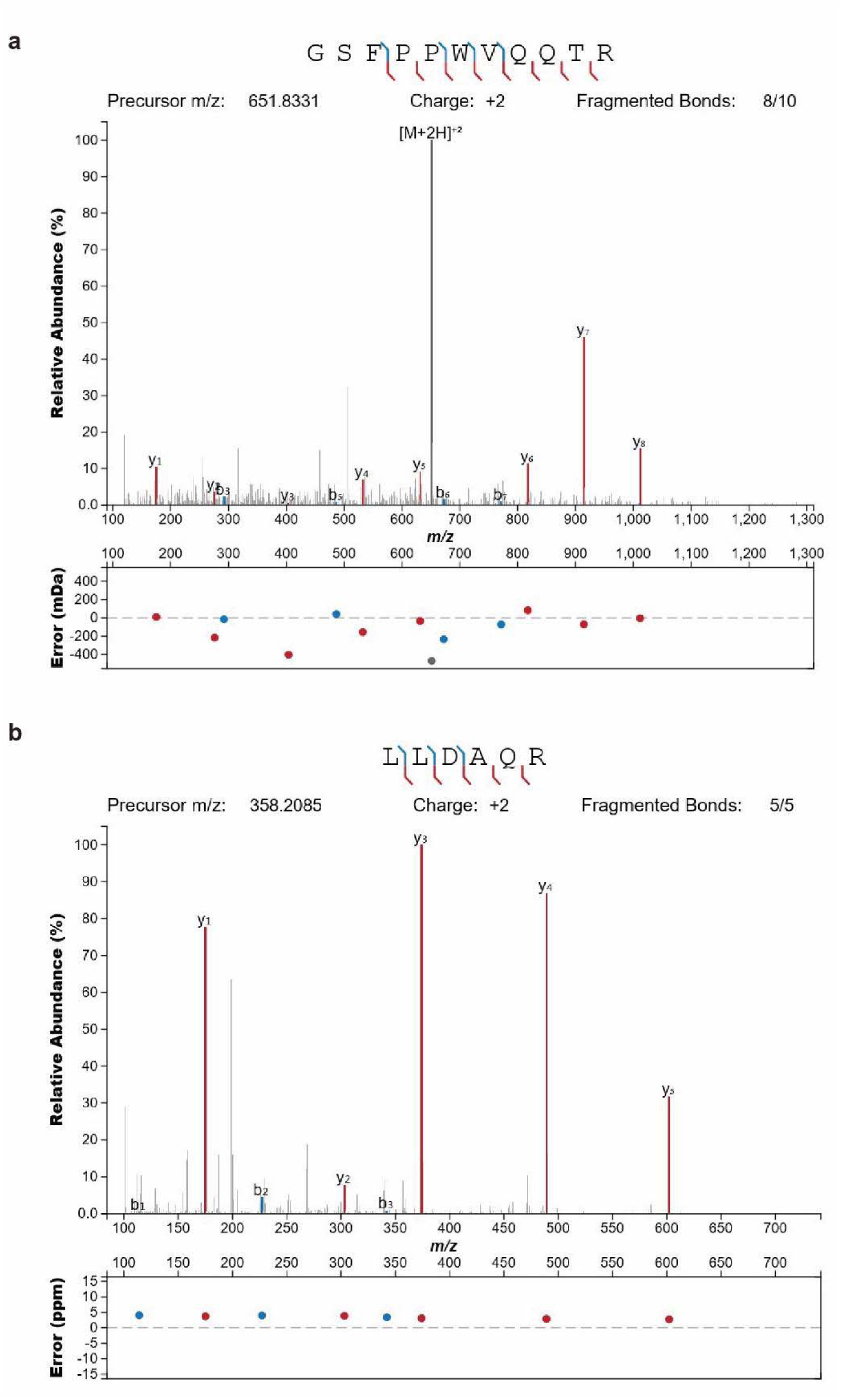
A. Annotated spectra of the SYNGAP1 isoform alpha 1 specific peptide “GSFPPWQQTR” identified from MS analysis of immune-isolated SYNGAP1 protein from D.I.V. 7 organoids. B. Annotated spectra of the SYNGAP1 isoform alpha 1 specific peptide “LLDAQR” identified from MS analysis of immune-isolated SYNGAP1 protein from D.I.V. 7 organoids.

**Supplementary Figure 3.**
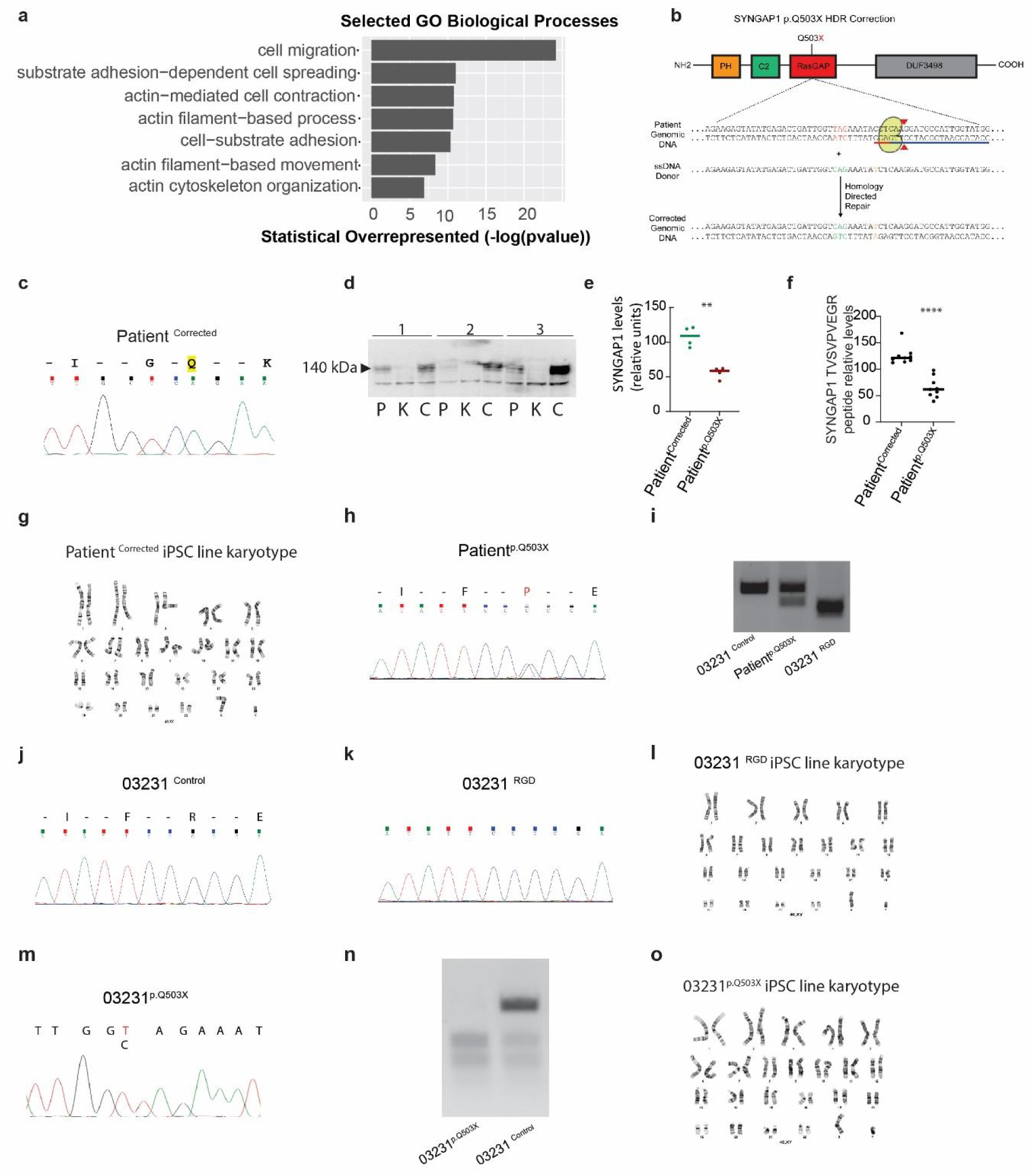
A. Selected GO terms for biological processes for SYNGAP1 immunoprecipitation data collected from D.I.V. 7 cortical organoids. B. Schematic of line generation details for isogenic control of Patient^p.Q503X^. C. Chromatogram of the generated corrected line (Patient^Corrected^). The truncating “T” was substituted with the wild type “C” base pair. D. Representative Western blot for SYNGAP1 in the Patient ^p.Q503X^(P), Patient^Corrected^ (C) and KO (K) iPSCs showing a reduction of SYNGAP1 levels in P and complete loss in K iPSCs E. Quantification of the western blot shows significant reduction in SYNGAP1 levels in Patient ^p.Q503X^ iPSCs compared to the Patient^Corrected^ iPSCs in four biological replicates. P<0.01.Individual dots represent independent replicates. Bar: mean values. Unpaired T-test. F. Quantification of the SYNGAP1 peptide. Graphical representation of SYNGAP1 protein levels quantified by timed parallel reaction monitoring (tRPM). Plot shows a decrease of SYNGAP1 total protein levels in Patient^p.Q503X^ :64.83 (51.6%) as compared to its corresponding isogenic control (Patient^Corrected^): 125.6 expressed as fg peptide/ug digested protein. N=9 across three independent differentiations. Unpaired t test P<0.0001 G. Karyotypic analysis of Patient^Corrected^ iPSCs revealed a normal karyotype. H. Chromatogram of the Patient^p.Q503X^ iPSCs carrying the truncating mutation. I. PCR of RGD, Patient^p.Q503X^, 03231 cell line. J. Chromatogram of the 03231^Control^ iPSCs carrying the wild type sequence K. Chromatogram of the 03231^RGD^ iPSCs carrying the homozygous mutation in the RGD domain. L. Karyotypic analysis of 03231^RGD^ iPSCs revealed a normal karyotype. M. Chromatogram of the 03231^p.Q503X^ iPSCs carrying the truncating mutation. N. PCR of 03231^Control^ cell line and 03231^p.Q503X^ cell line showing haploinsuffiency in the 03231^p.Q503X^ line. O. Karyotypic analysis of 03231^p.Q503X^ iPSCs revealed a normal karyotype.

**Supplementary Figure 4.**
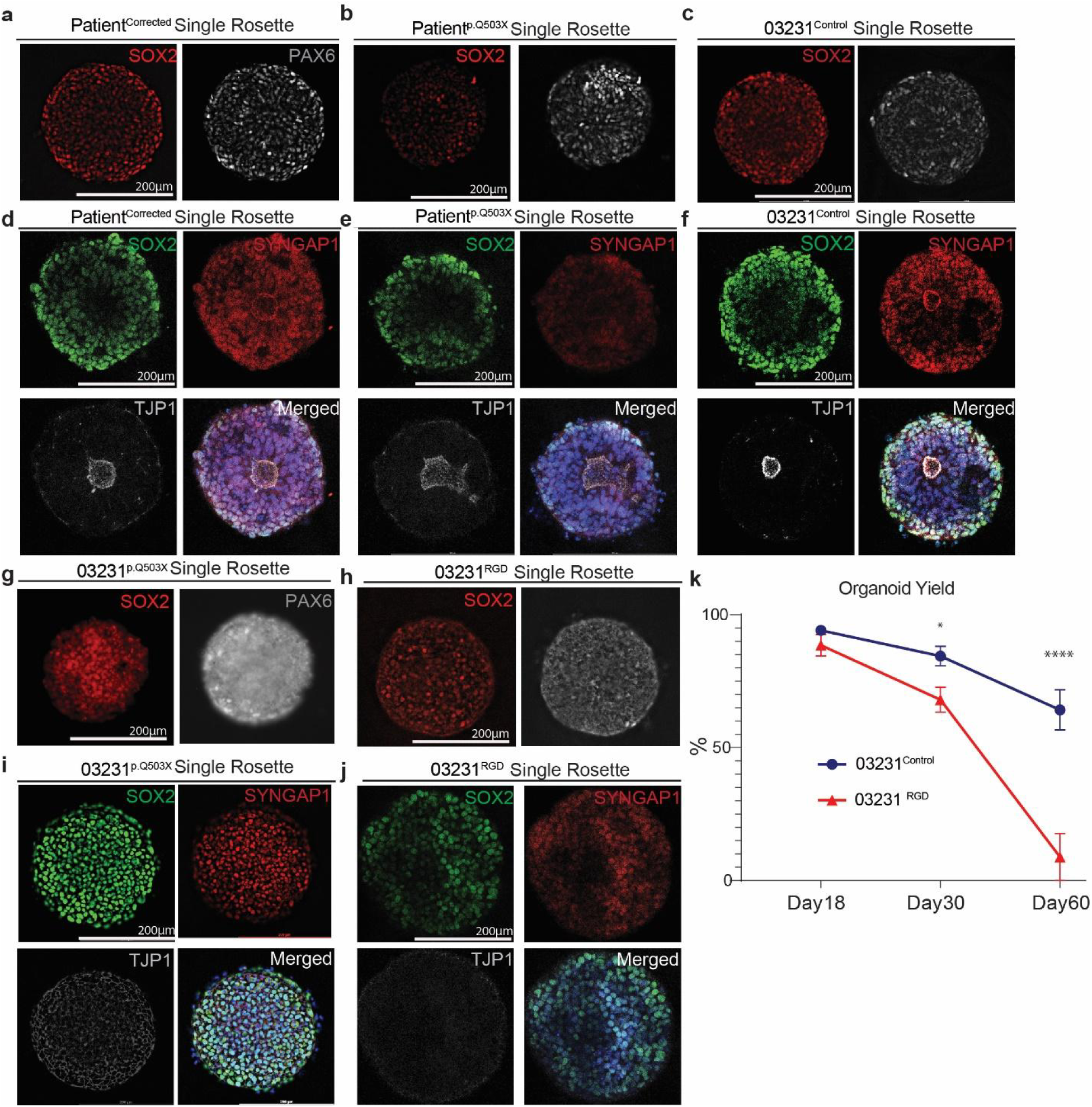
A. Single rosette from the Patient^Corrected^ cell line expressing SOX2 and PAX6 markers. B. Single rosette from the Patient^p.Q503X^ cell line expressing SOX2 and PAX6 markers. C. Single rosette from the 03231^Control^ cell line expressing SOX2 and PAX6 markers. D. A single rosette was generated from Patient^Corrected^ line. The rosette is composed of cells positive for the neural progenitor marker SOX2 and SYNGAP1. SYNGAP1 is also highly expressed at the apical wall of the lumen. The tight junction protein TJP1 labels the central luminal space of the rosette. Merged images show colocalization of DAPI, SYNGAP1, and TJP1. E. A single rosette was generated from Patient^p.Q503X^ line. The rosette is composed of cells positive for the neural progenitor marker SOX2 and SYNGAP1. Merged images show colocalization of DAPI, SYNGAP1, and TJP1. The Patient^p.Q503X^ single rosettes display a larger and more irregularly shaped central luminal space. F. A single rosette was generated from 03231^Control^ line. The rosette is composed of cells positive for the neural progenitor marker SOX2 and SYNGAP1. SYNGAP1 is also highly expressed at the apical wall of the lumen. Merged images show colocalization of DAPI, SYNGAP1, and TJP1. G. Single rosette from the 03231^p.Q503X^ cell line expressing SOX2 and PAX6 markers. H. Single rosette from the 03231^RGD^ cell line expressing SOX2 and PAX6 markers. I. A single rosette was generated from 03231^p.Q503X^ line. The rosette is composed of cells positive for the neural progenitor marker SOX2 and SYNGAP1. Merged images show colocalization of DAPI, SYNGAP1, and TJP1. The tight junction protein TJP1 is weakly expressed with little to no central luminal organization. J. A single rosette was generated from 03231^RGD^ line. The rosette is composed of cells positive for the neural progenitor marker SOX2 and SYNGAP1. Merged images show colocalization of DAPI, SYNGAP1, and TJP1. The tight junction protein TJP1 is weakly expressed with no central luminal organization. K. A Survival curve for organoids generated from the 03231^RGD^ line. Data was collected from 10 independent differentiations, each representing an average of 6 organoids for each time point. Single dots represent total averages for that time point. Student’s t-test was performed (Day 30 P=0.0134, Day 60 P<0.0001). Data is shown as mean ± SEM.

**Supplementary Figure 5.**
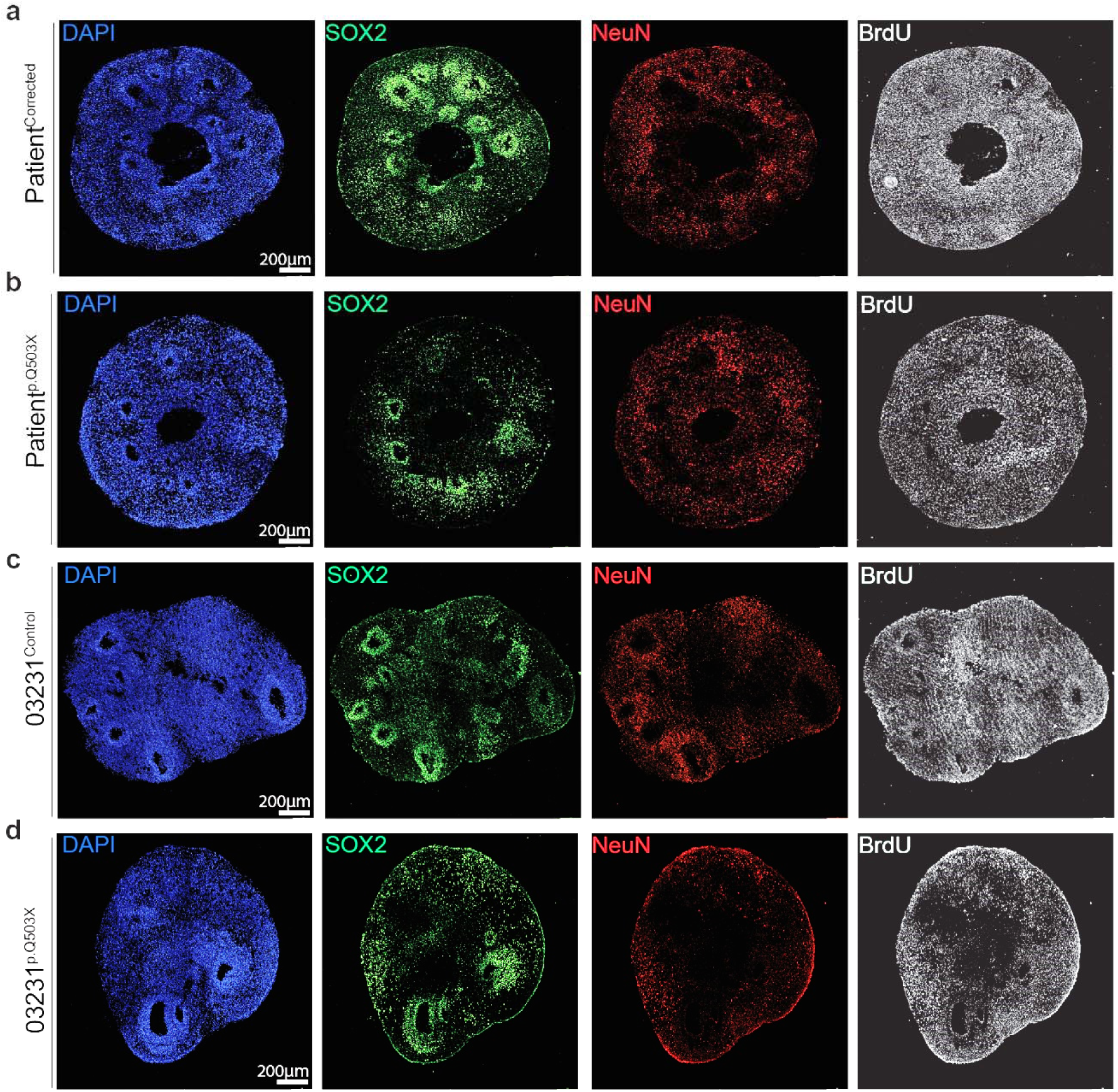
A. Representative single channel images from BrdU pulse-chase experiments in 2-month-old Patient^Corrected^ organoids. Images show the expression of the progenitor marker SOX2, the neuronal marker NeuN and the proliferative marker BrdU. B. Representative single channel images from BrdU pulse-chase experiments in 2-month-old Patient^p.Q503X^ organoids. Images show the expression of the progenitor marker SOX2, the neuronal marker NeuN and the proliferative marker BrdU. C. Representative single channel images from BrdU pulse-chase experiments in 2-month-old 03231^Control^ organoids. Images show the expression of the progenitor marker SOX2, the neuronal marker NeuN and the proliferative marker BrdU. D. Representative single channel images from BrdU pulse-chase experiments in 2-month-old 03231^p.Q503X^ organoids. Images show the expression of the progenitor marker SOX2, the neuronal marker NeuN and the proliferative marker BrdU.

**Supplementary Figure 6.**
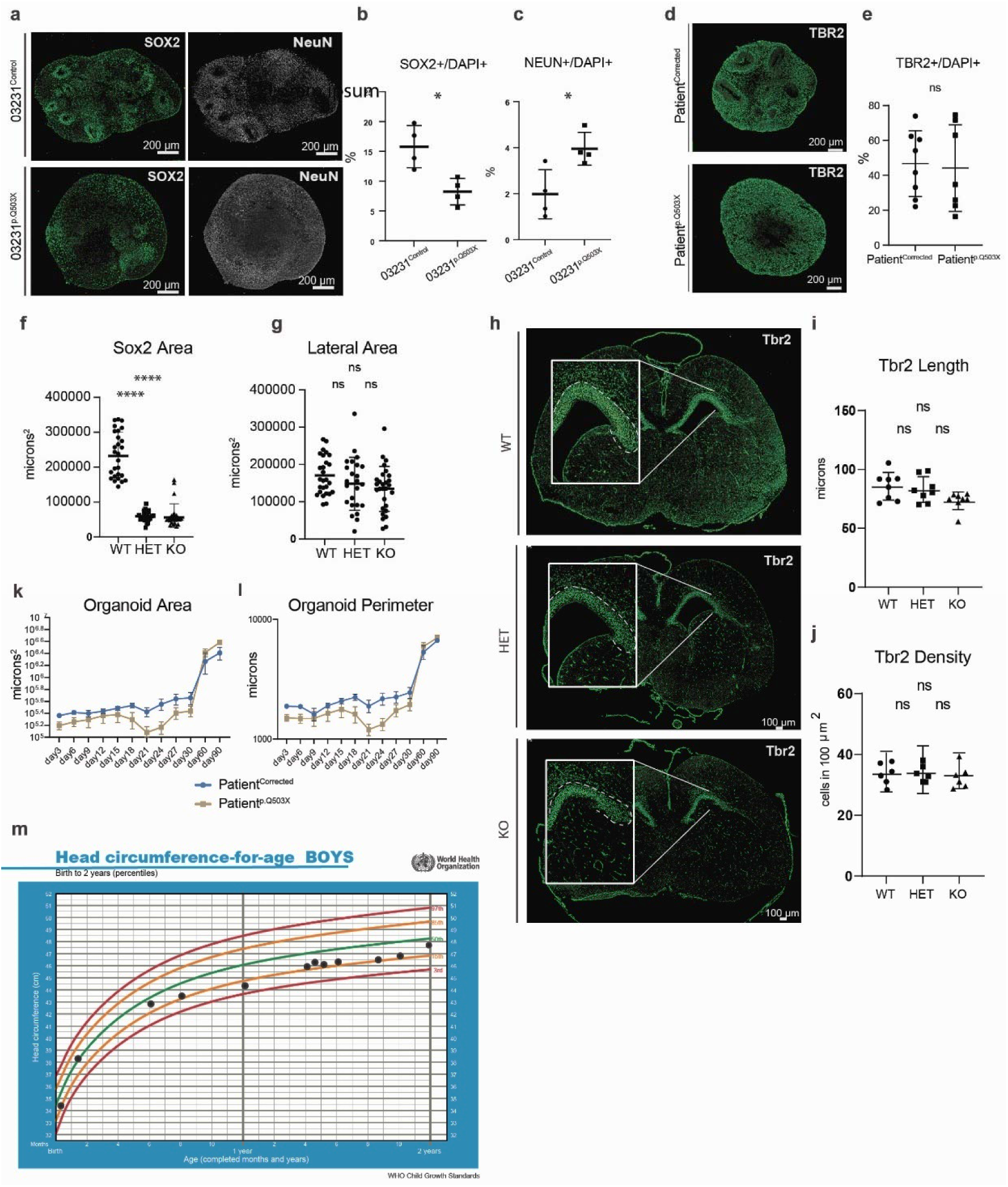
A. Representative images of SOX2 and NeuN expression in 2-month-old 03231^Control^ and 03231^p.Q503X^ organoids. B. Quantification of the total number of SOX2 positive cells normalized to DAP1. 03231^p.Q503X^ organoids show a reduced density of SOX2 positive progenitors indicating and earlier depletion of the progenitor pool. A Student’s t-test was performed on. Each dot represents an average value for all organoids from 1 differentiation. A Student’s t-test was performed on average values for 6-10 organoids from 4 differentiations. P Value =0.0112. Data is shown as mean ± SD. C. Quantification of the total number of NeuN positive cells normalized to DAP1. The higher density of NeuN positive cells in 03231^p.Q503X^ organoids indicates an accelerated differentiation of radial glial progenitors. A Student’s t-test was performed on average values for 6-10 organoids from 4 differentiations. P Value =0.0217. Data is shown as mean ± SD. D. Representative images of TBR2 expression in 2-month-old Patient^Corrected^ and Patient^p.Q503X^ organoids. E. Quantification of the total number of TBR2 positive intermediate progenitor cells normalized to DAP1 in 2-month-old Patient^Corrected^ and Patient^p.Q503X^ organoids showing no difference in TBR2 density between the lines. A Student’s t-test was performed on average values for 6-10 organoids from 4 differentiations. P= ns. Data is shown as mean ± SD. F. Quantification of the SOX2 positive area in the dorsal cortex of E18.5 mouse brains. Data was collected from 26-29 ventricles from 4 brains for each genotype. A One Way ANOVA was preformed between the genotypes, showing a decrease in the SOX2 positive progenitor regions in Het and KO as compared to WT mice. P<0.0001. Data is shown as mean ± SD. G. Quantification of the SOX2 positive area in the lateral cortex of E18.5 mouse brains. Data was collected from 26-29 ventricles from 4 brains for each genotype. A One Way ANOVA was preformed between the genotypes, showing no change in the SOX2 positive progenitor regions in Het and KO as compared to WT mice. P=ns. Data is shown as mean ± SD. H. Representative images of TBR2 expression in WT, Het and KO E18.5 mouse brains. I. Quantification of TBR2 thickness in the cortical plate in sections of E18.5 mouse brains. No change in the TBR2 thickness was observed between genotypes. A One Way ANOVA was performed on 8 ventricles from four animals for each genotype. P=ns. J. Quantification of the number of TBR2^+^ cells in 100 um^2^ of the VZ area. A One Way ANOVA was performed on 8-12 ventricles from four animals for each genotype. P=ns. K. Analysis of organoid area over time assessed by brightfield images for organoids generated from the Patient^Corrected^ and Patient^p.Q503X^ lines showing similar organoid size throughout long term culture. Data was collected from 10 independent differentiations, each representing an average of 6 organoids for each time point. Data is shown as mean ± SEM. L. Analysis of organoid perimeter over time assessed by brightfield images for organoids generated from the Patient^Corrected^ and Patient^p.Q503X^ lines showing similar organoid size throughout long term culture. Data was collected from 10 independent differentiations, each representing an average of 6 organoids for each time point. Data is shown as mean ± SEM. M. Head circumference measurements over time represented as dots from the Patient^p.Q503X^ donor plotted against the WHO child growth standards showing a consistent placing between the 3^rd^ and 15^th^ percentile.

**Supplementary Figure 7.**
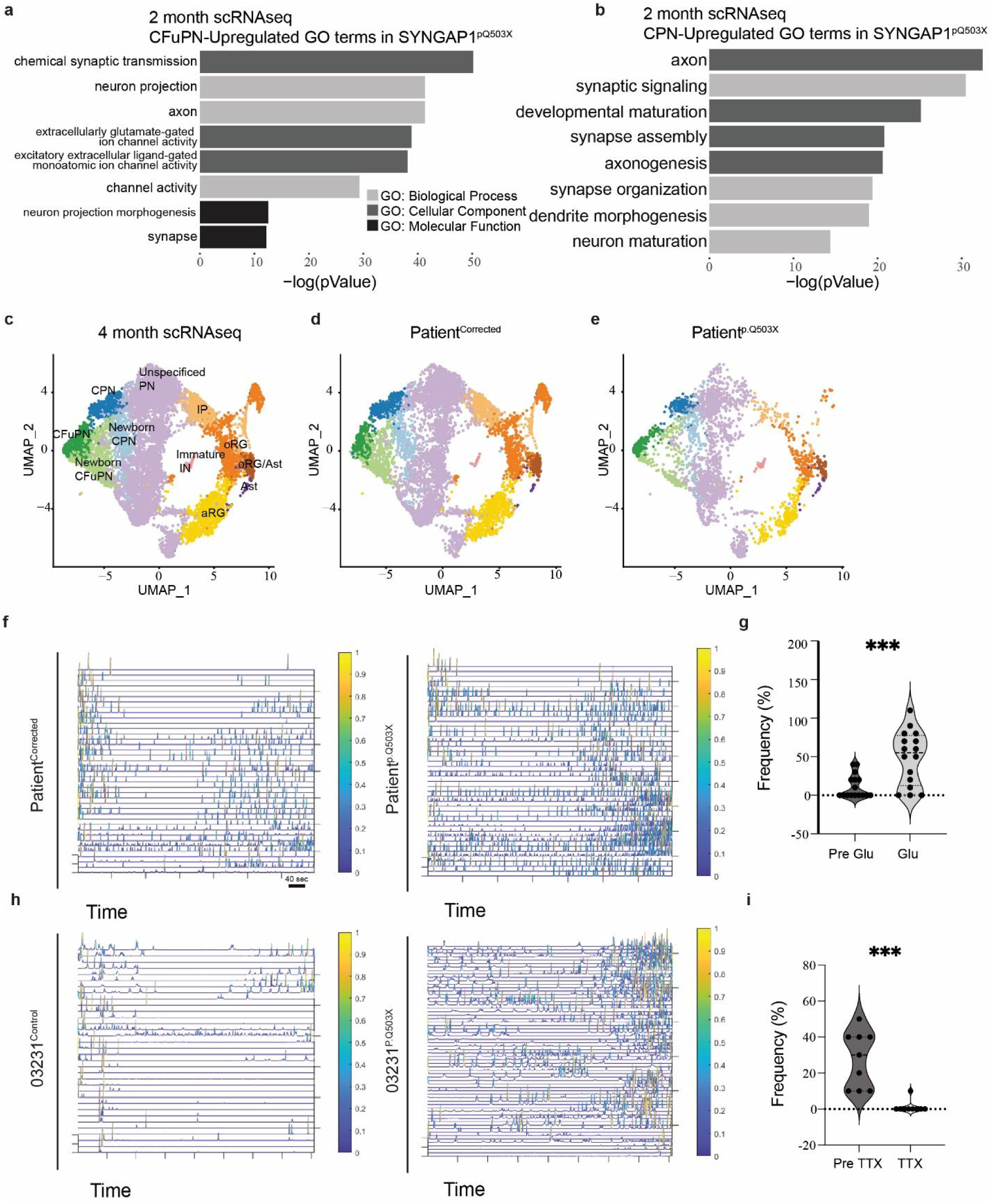
A. GO-Terms from single cell RNA sequencing preformed in 2-month-old Patient^p.Q503X^ and Patient^Corrected^ organoids. Graphical representation of upregulated terms for Patient^p.Q503X^ Corticofugal Projection Neurons (CFuPN). Main biological process, cellular component, and molecular function GO-Terms are related to neuronal differentiation and synapse formation. B. GO-Terms from single cell RNA sequencing preformed in 4-month-old Patient^p.Q503X^ and Patient^Corrected^ organoids. Graphical representation of upregulated GO-Terms in Patient^p.Q503X^ Callosal Projection Neurons (CPN). Main biological process and cellular component GO-Terms are related to neuronal differentiation and synapse formation. C. Combined t-distributed stochastic neighbor embedding (t-SNE) from single cell RNA sequencing analysis of pooled Patientp.^Q503X^ and Patient^Corrected^ organoids at 4 months. D. Individual t-SNE plot for pooled Patient^corrected^ organoids at 4 months (n=7540 cells). E. Individual t-SNE plot for pooled Patient^p.Q503X^ organoids at 4 months (n=3123 cells). F. ΔF/F(t) from GCaMP6f2 recordings of Patient^Corrected^ and Patient^p.Q503X^ organoids. G. Calcium spike frequency analysis on 2-month-old Patient^p.Q503X^ organoids before and during bath application of glutamate (Glu). Unpaired t-test performed on a total of 16 cells from Patient^p.Q503X^ organoids, from 3 independent experiments. P value = 0.0005. Data is shown as mean ± SD H. ΔF/F(t) from GCaMP6f2 recordings of 03231^Control^ and 03231^p.Q503X^ organoids. I. Calcium spike frequency analysis on 2-month-old Patient^p.Q503X^ organoids before and during bath application of tetrodotoxin (TTX). Unpaired t-test performed on a total of 9 cells from Patient^p.Q503X^ organoids, from 3 independent experiments. P value =0.0001. Data is shown as mean ± SD

**Supplementary Table 1**

Proteome of 7 D.I.V. Cortical Organoids.

**Supplementary Table 2**

SynGO analysis of 7 D.I.V. Cortical Organoids Proteome.

**Supplementary Table 3**

Interactome of SYNGAP1 in 7 D.I.V. Cortical Organoids.

**Supplementary Table 4**

Off-Target Sequences and Sequencing.

**SYNGAP1 p.Q503X-Corrected gRNA Predicted Off-Target Sequences and Sequencing**. Table contains the top predicted off-target regions of the guide RNA used to generate Patient 1-Corrected iPSCs along with sequencing results of each target that was PCR amplified and then verified via Sanger Sequencing. The predicted off-target sequence is highlighted along with 25 base pairs on either side.

**RASGAP dead gRNA Predicted Off-Target Sequences and Sequencing**. Table contains the top ten predicted off-target regions of the guide RNA used to generate RASGAP dead iPSCs along with sequencing results of each target that was PCR amplified and then verified via Sanger Sequencing. The predicted off-target sequence is highlighted along with 25 base pairs on either side.

**Supplementary Table 5**

SYNGAP1 quantitation

Table shows MS methods, acceptance criteria, peptides concentrations, calibration curve and statistical analysis for PRM quantitation of total protein levels of SYNGAP1

**Supplementary Table 6**

Differential Expression Analysis (DEGs) from Bulk-RNA sequencing of 7 D.I.V. Cortical Organoids from SYNGAP1^p.Q503X^ and corrected line.

**Supplementary Table 7**

Differential Expression Analysis (DEGs) from SC-RNA sequencing of 2-month-old Cortical Organoids from SYNGAP1^p.Q503X^ and corrected line.

**Supplementary Table 8**

Differential Expression Analysis (DEGs) from SC-RNA sequencing of 4-month-old Cortical Organoids from SYNGAP1^p.Q503X^ and corrected line.

**Supplementary Video 1**

Live imaging of corrected rosette formation from day 5 to day 7 after initial seeding.

**Supplementary Video 2**

Live imaging of SYNGAP1^p.Q503X^ rosette formation from day 5 to day 7 after initial seeding.

**Supplementary Video 3**

10 minutes video recordings of GCaMP6f corrected organoids.

**Supplementary Video 4**

10 minutes video recordings of GCaMP6f SYNGAP1^p.Q503X^ organoids.

**Supplementary Video 5**

10 minutes video recordings of GCaMP6f 03231 organoids.

**Supplementary Video 6**

10 minutes video recordings of GCaMP6f 03231^p.Q503X^ organoids.

## References

1 De Rubeis, S. et al. Synaptic, transcriptional and chromatin genes disrupted in autism. Nature 515, 209–215 (2014). 10.1038/nature13772

2 Paulsen, B. et al. Autism genes converge on asynchronous development of shared neuron classes. Nature 602, 268–273 (2022). 10.1038/s41586-021-04358-6

3 Mariani, J. et al. FOXG1-Dependent Dysregulation of GABA/Glutamate Neuron Differentiation in Autism Spectrum Disorders. Cell 162, 375–390 (2015). 10.1016/j.cell.2015.06.034

4 Jourdon, A. et al. ASD modelling in organoids reveals imbalance of excitatory cortical neuron subtypes during early neurogenesis. bioRxiv, 2022.2003.2019.484988 (2022). 10.1101/2022.03.19.484988

5 Villa, C. E. et al. CHD8 haploinsufficiency links autism to transient alterations in excitatory and inhibitory trajectories. Cell Rep 39, 110615 (2022). 10.1016/j.celrep.2022.110615

6 Schafer, S. T. et al. Pathological priming causes developmental gene network heterochronicity in autistic subject-derived neurons. Nat Neurosci 22, 243–255 (2019). 10.1038/s41593-018-0295-x

7 de Jong, J. O. et al. Cortical overgrowth in a preclinical forebrain organoid model of CNTNAP2-associated autism spectrum disorder. Nat Commun 12, 4087 (2021). 10.1038/s41467-021-24358-4

8 Urresti, J. et al. Cortical organoids model early brain development disrupted by 16p11.2 copy number variants in autism. Mol Psychiatry 26, 7560–7580 (2021). 10.1038/s41380-021-01243-6

9 Satterstrom, F. K. et al. Large-Scale Exome Sequencing Study Implicates Both Developmental and Functional Changes in the Neurobiology of Autism. Cell 180, 568–584.e523 (2020). 10.1016/j.cell.2019.12.036

10 Chen, H. J., Rojas-Soto, M., Oguni, A. & Kennedy, M. B. A synaptic Ras-GTPase activating protein (p135 SynGAP) inhibited by CaM kinase II. Neuron 20, 895–904 (1998). 10.1016/s0896-6273(00)80471-7

11 Kim, J. H., Liao, D., Lau, L. F. & Huganir, R. L. SynGAP: a synaptic RasGAP that associates with the PSD-95/SAP90 protein family. Neuron 20, 683–691 (1998).

12 Komiyama, N. H. et al. SynGAP regulates ERK/MAPK signaling, synaptic plasticity, and learning in the complex with postsynaptic density 95 and NMDA receptor. J Neurosci 22, 9721–9732 (2002).

13 Kim, J. H., Lee, H. K., Takamiya, K. & Huganir, R. L. The role of synaptic GTPase-activating protein in neuronal development and synaptic plasticity. J Neurosci 23, 1119–1124 (2003).

14 Zhu, J. J., Qin, Y., Zhao, M., Van Aelst, L. & Malinow, R. Ras and Rap control AMPA receptor trafficking during synaptic plasticity. Cell 110, 443–455 (2002). 10.1016/s0092-8674(02)00897-8

15 Araki, Y., Zeng, M., Zhang, M. & Huganir, R. L. Rapid dispersion of SynGAP from synaptic spines triggers AMPA receptor insertion and spine enlargement during LTP. Neuron 85, 173–189 (2015). 10.1016/j.neuron.2014.12.023

16 Walkup, W. G. et al. A model for regulation by SynGAP-α1 of binding of synaptic proteins to PDZ-domain ‘Slots’ in the postsynaptic density. Elife 5 (2016). 10.7554/eLife.16813

17 Zeng, M. et al. Phase Transition in Postsynaptic Densities Underlies Formation of Synaptic Complexes and Synaptic Plasticity. Cell 166, 1163–1175.e1112 (2016). 10.1016/j.cell.2016.07.008

18 Zeng, M., Bai, G. & Zhang, M. Anchoring high concentrations of SynGAP at postsynaptic densities via liquid-liquid phase separation. Small GTPases 10, 296–304 (2019). 10.1080/21541248.2017.1320350

19 Kilinc, M. et al. Endogenous. Elife 11 (2022). 10.7554/eLife.75707

20 Knuesel, I., Elliott, A., Chen, H. J., Mansuy, I. M. & Kennedy, M. B. A role for synGAP in regulating neuronal apoptosis. Eur J Neurosci 21, 611–621 (2005). 10.1111/j.1460-9568.2005.03908.x

21 Willsey, H. R. et al. Parallel in vivo analysis of large-effect autism genes implicates cortical neurogenesis and estrogen in risk and resilience. Neuron 109, 1409 (2021). 10.1016/j.neuron.2021.03.030

22 Su, P. et al. Disruption of SynGAP-dopamine D1 receptor complexes alters actin and microtubule dynamics and impairs GABAergic interneuron migration. Sci Signal 12 (2019). 10.1126/scisignal.aau9122

23 Berryer, M. H. et al. Mutations in SYNGAP1 cause intellectual disability, autism, and a specific form of epilepsy by inducing haploinsufficiency. Hum Mutat 34, 385–394 (2013). 10.1002/humu.22248

24 Kilinc, M. et al. Species-conserved SYNGAP1 phenotypes associated with neurodevelopmental disorders. Mol Cell Neurosci 91, 140–150 (2018). 10.1016/j.mcn.2018.03.008

25 Gamache, T. R., Araki, Y. & Huganir, R. L. Twenty Years of SynGAP Research: From Synapses to Cognition. The Journal of Neuroscience 40, 1596 (2020). 10.1523/JNEUROSCI.0420-19.2020

26 Lancaster, M. A. et al. Cerebral organoids model human brain development and microcephaly. Nature 501, 373–379 (2013). 10.1038/nature12517

27 Bershteyn, M. et al. Human iPSC-Derived Cerebral Organoids Model Cellular Features of Lissencephaly and Reveal Prolonged Mitosis of Outer Radial Glia. Cell Stem Cell 20, 435–449.e434 (2017). 10.1016/j.stem.2016.12.007

28 Kanton, S. et al. Organoid single-cell genomic atlas uncovers human-specific features of brain development. Nature 574, 418–422 (2019). 10.1038/s41586-019-1654-9

29 Klaus, J. et al. Altered neuronal migratory trajectories in human cerebral organoids derived from individuals with neuronal heterotopia. Nat Med 25, 561–568 (2019). 10.1038/s41591-019-0371-0

30 Esk, C. et al. A human tissue screen identifies a regulator of ER secretion as a brain-size determinant. Science 370, 935–941 (2020). 10.1126/science.abb5390

31 Khan, T. A. et al. Neuronal defects in a human cellular model of 22q11.2 deletion syndrome. Nat Med 26, 1888–1898 (2020). 10.1038/s41591-020-1043-9

32 Samarasinghe, R. A. et al. Identification of neural oscillations and epileptiform changes in human brain organoids. Nat Neurosci 24, 1488–1500 (2021). 10.1038/s41593-021-00906-5

33 Tidball, A. M. et al. Self-organizing Single-Rosette Brain Organoids from Human Pluripotent Stem Cells. bioRxiv, 2022.2002.2028.482350 (2022). 10.1101/2022.02.28.482350

34 Quadrato, G. et al. Cell diversity and network dynamics in photosensitive human brain organoids. Nature 545, 48–53 (2017). 10.1038/nature22047

35 Velasco, S. et al. Individual brain organoids reproducibly form cell diversity of the human cerebral cortex. Nature 570, 523–527 (2019). 10.1038/s41586-019-1289-x

36 Camp, J. G. et al. Human cerebral organoids recapitulate gene expression programs of fetal neocortex development. Proc Natl Acad Sci U S A 112, 15672–15677 (2015). 10.1073/pnas.1520760112

37 Trevino, A. E. et al. Chromatin accessibility dynamics in a model of human forebrain development. Science 367 (2020). 10.1126/science.aay1645

38 Gordon, A. et al. Long-term maturation of human cortical organoids matches key early postnatal transitions. Nat Neurosci 24, 331–342 (2021). 10.1038/s41593-021-00802-y

39 Michaelson, S. D. et al. SYNGAP1 heterozygosity disrupts sensory processing by reducing touch-related activity within somatosensory cortex circuits. Nat Neurosci 21, 1–13 (2018). 10.1038/s41593-018-0268-0

40 Aceti, M. et al. Syngap1 haploinsufficiency damages a postnatal critical period of pyramidal cell structural maturation linked to cortical circuit assembly. Biol Psychiatry 77, 805–815 (2015). 10.1016/j.biopsych.2014.08.001

41 Eze, U. C., Bhaduri, A., Haeussler, M., Nowakowski, T. J. & Kriegstein, A. R. Single-cell atlas of early human brain development highlights heterogeneity of human neuroepithelial cells and early radial glia. Nat Neurosci 24, 584–594 (2021). 10.1038/s41593-020-00794-1

42 Li, J. et al. Spatiotemporal profile of postsynaptic interactomes integrates components of complex brain disorders. Nat Neurosci 20, 1150–1161 (2017). 10.1038/nn.4594

43 Araki, Y. et al. SynGAP isoforms differentially regulate synaptic plasticity and dendritic development. Elife 9 (2020). 10.7554/eLife.56273

44 Knight, G. T. et al. Engineering induction of singular neural rosette emergence within hPSC-derived tissues. Elife 7 (2018). 10.7554/eLife.37549

45 Wilkinson, B. et al. Endogenous Cell Type-Specific Disrupted in Schizophrenia 1 Interactomes Reveal Protein Networks Associated With Neurodevelopmental Disorders. Biol Psychiatry 85, 305–316 (2019). 10.1016/j.biopsych.2018.05.009

46 Carlisle, H. J., Manzerra, P., Marcora, E. & Kennedy, M. B. SynGAP regulates steady-state and activity-dependent phosphorylation of cofilin. J Neurosci 28, 13673–13683 (2008). 10.1523/JNEUROSCI.4695-08.2008

47 Tomoda, T., Kim, J. H., Zhan, C. & Hatten, M. E. Role of Unc51.1 and its binding partners in CNS axon outgrowth. Genes Dev 18, 541–558 (2004). 10.1101/gad.1151204

48 Aaku-Saraste, E., Hellwig, A. & Huttner, W. B. Loss of occludin and functional tight junctions, but not ZO-1, during neural tube closure--remodeling of the neuroepithelium prior to neurogenesis. Dev Biol 180, 664–679 (1996). 10.1006/dbio.1996.0336

49 Edmondson, J. C. & Hatten, M. E. Glial-guided granule neuron migration in vitro: a high-resolution time-lapse video microscopic study. J Neurosci 7, 1928–1934 (1987).

50 Rakic, P. Guidance of neurons migrating to the fetal monkey neocortex. Brain Res 33, 471–476 (1971). 10.1016/0006-8993(71)90119-3

51 Rakic, P. Mode of cell migration to the superficial layers of fetal monkey neocortex. J Comp Neurol 145, 61–83 (1972). 10.1002/cne.901450105

52 Rakic, P. Neuronal migration and contact guidance in the primate telencephalon. Postgrad Med J 54 **Suppl 1**, 25–40 (1978).

53 Nowakowski, T. J., Pollen, A. A., Sandoval-Espinosa, C. & Kriegstein, A. R. Transformation of the Radial Glia Scaffold Demarcates Two Stages of Human Cerebral Cortex Development. Neuron 91, 1219–1227 (2016). 10.1016/j.neuron.2016.09.005

54 Chenn, A. & McConnell, S. K. Cleavage orientation and the asymmetric inheritance of Notch1 immunoreactivity in mammalian neurogenesis. Cell 82, 631–641 (1995). 10.1016/0092-8674(95)90035-7

55 Shitamukai, A., Konno, D. & Matsuzaki, F. Oblique radial glial divisions in the developing mouse neocortex induce self-renewing progenitors outside the germinal zone that resemble primate outer subventricular zone progenitors. J Neurosci 31, 3683–3695 (2011). 10.1523/JNEUROSCI.4773-10.2011

56 Nowakowski, T. J. et al. Spatiotemporal gene expression trajectories reveal developmental hierarchies of the human cortex. Science 358, 1318–1323 (2017). 10.1126/science.aap8809

57 Llamosas, N. et al. SYNGAP1 Controls the Maturation of Dendrites, Synaptic Function, and Network Activity in Developing Human Neurons.

58 Fromer, M. et al. De novo mutations in schizophrenia implicate synaptic networks. Nature 506, 179–184 (2014). 10.1038/nature12929

59 Genovese, G. et al. Increased burden of ultra-rare protein-altering variants among 4,877 individuals with schizophrenia. Nat Neurosci 19, 1433–1441 (2016). 10.1038/nn.4402

60 Kirov, G. et al. De novo CNV analysis implicates specific abnormalities of postsynaptic signalling complexes in the pathogenesis of schizophrenia. Mol Psychiatry 17, 142–153 (2012). 10.1038/mp.2011.154

61 O’Roak, B. J. et al. Exome sequencing in sporadic autism spectrum disorders identifies severe de novo mutations. Nat Genet 43, 585–589 (2011). 10.1038/ng.835

62 O’Roak, B. J. et al. Sporadic autism exomes reveal a highly interconnected protein network of de novo mutations. Nature 485, 246–250 (2012). 10.1038/nature10989

63 Peça, J. & Feng, G. Cellular and synaptic network defects in autism. Curr Opin Neurobiol 22, 866–872 (2012). 10.1016/j.conb.2012.02.015

64 Kawaguchi, A. Neuronal Delamination and Outer Radial Glia Generation in Neocortical Development. Frontiers in Cell and Developmental Biology 8 (2021).

65 Kadowaki, M. et al. N-cadherin mediates cortical organization in the mouse brain. Dev Biol 304, 22–33 (2007). 10.1016/j.ydbio.2006.12.014

66 Cappello, S. et al. A radial glia-specific role of RhoA in double cortex formation. Neuron 73, 911–924 (2012). 10.1016/j.neuron.2011.12.030

67 Gil-Sanz, C., Landeira, B., Ramos, C., Costa, M. R. & Müller, U. Proliferative defects and formation of a double cortex in mice lacking Mltt4 and Cdh2 in the dorsal telencephalon. J Neurosci 34, 10475–10487 (2014). 10.1523/JNEUROSCI.1793-14.2014

68 Yoon, K. J. et al. Modeling a genetic risk for schizophrenia in iPSCs and mice reveals neural stem cell deficits associated with adherens junctions and polarity. Cell Stem Cell 15, 79–91 (2014). 10.1016/j.stem.2014.05.003

69 Hansen, D. V., Lui, J. H., Parker, P. R. & Kriegstein, A. R. Neurogenic radial glia in the outer subventricular zone of human neocortex. Nature 464, 554–561 (2010). 10.1038/nature08845

70 Clement, J. P., Ozkan, E. D., Aceti, M., Miller, C. A. & Rumbaugh, G. SYNGAP1 links the maturation rate of excitatory synapses to the duration of critical-period synaptic plasticity. J Neurosci 33, 10447–10452 (2013). 10.1523/JNEUROSCI.0765-13.2013

71 Clement, J. P. et al. Pathogenic SYNGAP1 mutations impair cognitive development by disrupting maturation of dendritic spine synapses. Cell 151, 709–723 (2012). 10.1016/j.cell.2012.08.045

72 Ozkan, E. D. et al. Reduced cognition in Syngap1 mutants is caused by isolated damage within developing forebrain excitatory neurons. Neuron 82, 1317–1333 (2014). 10.1016/j.neuron.2014.05.015

73 Creson, T. K. et al. Re-expression of SynGAP protein in adulthood improves translatable measures of brain function and behavior. Elife 8 (2019). 10.7554/eLife.46752

74 Noctor, S. C., Martínez-Cerdeño, V., Ivic, L. & Kriegstein, A. R. Cortical neurons arise in symmetric and asymmetric division zones and migrate through specific phases. Nat Neurosci 7, 136–144 (2004). 10.1038/nn1172

75 He, S., Li, Z., Ge, S., Yu, Y. C. & Shi, S. H. Inside-Out Radial Migration Facilitates Lineage-Dependent Neocortical Microcircuit Assembly. Neuron 86, 1159–1166 (2015). 10.1016/j.neuron.2015.05.002

76 Coba, M. P. et al. Dlgap1 knockout mice exhibit alterations of the postsynaptic density and selective reductions in sociability. Sci Rep 8, 2281 (2018). 10.1038/s41598-018-20610-y

77 Li, J. et al. Long-term potentiation modulates synaptic phosphorylation networks and reshapes the structure of the postsynaptic interactome. Sci Signal 9, rs8 (2016). 10.1126/scisignal.aaf6716

78 Romero, D. M. et al. Novel role of the synaptic scaffold protein Dlgap4 in ventricular surface integrity and neuronal migration during cortical development. Nat Commun 13, 2746 (2022). 10.1038/s41467-022-30443-z

79 Okita, K. et al. An efficient nonviral method to generate integration-free human-induced pluripotent stem cells from cord blood and peripheral blood cells. Stem Cells 31, 458–466 (2013). 10.1002/stem.1293

80 Klose, A. et al. Selective disactivation of neurofibromin GAP activity in neurofibromatosis type 1. Hum Mol Genet 7, 1261–1268 (1998). 10.1093/hmg/7.8.1261

81 Concordet, J. P. & Haeussler, M. CRISPOR: intuitive guide selection for CRISPR/Cas9 genome editing experiments and screens. Nucleic Acids Res 46, W242–W245 (2018). 10.1093/nar/gky354

82 Lippmann, E. S., Estevez-Silva, M. C. & Ashton, R. S. Defined human pluripotent stem cell culture enables highly efficient neuroepithelium derivation without small molecule inhibitors. Stem Cells 32, 1032–1042 (2014). 10.1002/stem.1622

83 Hughes, C. S. et al. Single-pot, solid-phase-enhanced sample preparation for proteomics experiments. Nat Protoc 14, 68–85 (2019). 10.1038/s41596-018-0082-x

84 Uzquiano, A. et al. Single-cell multiomics atlas of organoid development uncovers longitudinal molecular programs of cellular diversification of the human cerebral cortex. bioRxiv, 2022.2003.2017.484798 (2022). 10.1101/2022.03.17.484798

85 Giovannucci, A. et al. CaImAn an open source tool for scalable calcium imaging data analysis. Elife 8 (2019). 10.7554/eLife.38173

